# Neuroprotective Effects of Naltrexone in a Mouse Model of Post-Traumatic Epilepsy

**DOI:** 10.1101/2023.10.04.560949

**Authors:** Saul Rodriguez, Shaunik Sharma, Grant Tiarks, Zeru Peterson, Kyle Jackson, Daniel Thedens, Angela Wong, David Keffala-Gerhard, Vinit B. Mahajan, Polly J. Ferguson, Elizabeth A. Newell, Joseph Glykys, Thomas Nickl-Jockschat, Alexander G. Bassuk

**Affiliations:** Department of Pediatrics, University of Iowa, Iowa City, IA; Department of Psychiatry, University of Iowa, Iowa City, IA; Department of Radiology, University of Iowa, Iowa City, IA; Iowa Neuroscience Institute, University of Iowa, Iowa City, IA; Department of Neurology, University of Iowa, Iowa City, IA; Department of Neuroscience and Pharmacology, University of Iowa, Iowa City, IA; Department of Ophthalmology, Stanford University, Palo Alto, California

**Keywords:** Mu-opioid receptors, naltrexone, traumatic brain injury, TBI, epileptogenesis, nitro-oxidative stress, neuroinflammation, inflammatory cytokines

## Abstract

Traumatic Brain Injury (TBI) induces neuroinflammatory responses that can initiate epileptogenesis, which develops into epilepsy. Recently, we identified the anti-convulsive effects of naltrexone, a mu-opioid receptor (MOR) antagonist. While blocking opioid receptors can reduce inflammation, it is unclear if post-TBI epileptogenesis can be prevented by blocking MORs. Here, we tested if naltrexone prevents neuroinflammation and epileptogenesis post-TBI. TBI was induced by a modified Marmarau Weight-Drop (WD) method applied to four-week-old C57BL/6J male mice. Mice were given the pro-convulsant pentylenetetrazol (PTZ) on day two post-injury while telemetry-monitored mice received PTZ on day five. Naltrexone/vehicle treatment started two hours after PTZ. Integrated EEG-video (vEEG) recorded interictal events and spontaneous seizures for three months. Molecular, histological and neuroimaging techniques were used to evaluate neuroinflammation and neurodegeneration both acutely and chronically. Peripheral immune responses were assessed through serum chemokine/cytokine measurements. We observed increases in MOR expression, nitro-oxidative stress, mRNA expression of inflammatory cytokines, microgliosis, neurodegeneration, and white matter damage in the neocortex of TBI mice. vEEG revealed increased interictal events in TBI mice, with 71% developing epilepsy. Naltrexone ameliorated neuroinflammation and neurodegeneration, reduced interictal events and prevented epilepsy, illustrating that naltrexone is a promising drug to prevent TBI-associated neuroinflammation and epileptogenesis in post-traumatic epilepsy.

## Introduction

Traumatic brain injuries (TBIs) trigger focal and diffuse injury, increasing the probability of developing post-traumatic seizures and Post-traumatic Epilepsy (PTE) (1–3). PTE is the most common form of acquired epilepsy, accounting for L20% of symptomatic epilepsies (2). PTE is a complex, broadly heterogeneous condition, and its pathogenic mechanisms in the context of TBI and other traumas are incompletely understood. A neuroinflammatory response immediately follows traumatic brain injury (TBI) and is mediated by cytokines, chemokines, and free radicals. TBI also promotes blood-brain barrier breakdown, further exacerbating inflammation (4, 5).

Opioid receptors, particularly mu-opioid receptors (MORs), regulate neuroinflammatory responses and cause inflammation when overstimulated. Studies have shown that the MOR agonist morphine activates the transcription factor NF-ĸB, which in turn mediates a substantial immune response in microglia, resulting in inflammation. In contrast, silencing MORs through siRNA reduces NF-ĸB activation and impairs glial activation and transcription of inflammatory genes (6). Overstimulation of MORs also drives redox signaling and facilitates activation of various transcription factors in microglia, astrocytes, and neurons, at least in part via mitogen-activated protein kinase (MAPK) signaling (7). In glial cells, the signaling crosstalk between MOR and MAPK enhances the release of inflammatory cytokines (e.g., TNFα, IL-1β, and IL-6) and of inducible nitric oxide synthase (iNOS), which jointly increase the oxidative load, damaging cellular components (8, 9). Conversely, antagonizing or deleting MORs decreases neuroinflammation by attenuating oxidative stress and reducing inflammatory cytokines. These findings indicate that MOR signaling contributes to inflammation (10–12).

Opioid receptors not only modulate neuroinflammation but also play a complex role in seizures and can generate either a pro- or anti-convulsive effect depending on the dose, the drug administered, and the opioid receptor activated. We recently reported that naltrexone, a MOR antagonist, decreases seizure-like activity in zebrafish and mice *in vitro* and *in vivo* (*13*). Nonetheless, studies evaluating naltrexone’s long-term anti-epileptic properties are still lacking. Here, we investigated the anti-inflammatory and anti-epileptic effects of naltrexone in a modified Marmarou weight-drop (WD) model of TBI. We hypothesize that an increase in MOR activity in the neocortex after TBI would drive neuroinflammation and neurodegeneration, promoting epileptogenesis. In contrast, we expect that antagonizing MOR soon after TBI should reduce the severity of these inflammatory and degenerative processes in the brain and prevent the development of epilepsy. Accordingly, we tested the neuroprotective effects of naltrexone in the acute and chronic phases following TBI. The onset and progression of epileptogenesis were tracked by integrated remote video-EEG and by profiling epileptogenic biomarkers. We found an increase in epileptogenic and neuroinflammatory biomarkers in the neocortex of TBI mice, whereas intervention with naltrexone after injury decreased those markers, prevented widespread injury to the white matter (as determined through diffusion tensor magnetic resonance imaging) and prevented epilepsy. Our results show that targeting MORs can provide a therapeutic advantage against the neuroinflammation which starts immediately after TBI and ultimately prevent the progression of epileptogenesis.

## Results

### Naltrexone decreased phosphorylation and expression of MOR in the neocortex

The time points for the TBI experiment, the PTZ test conditions, and the protocol for electrode implantation for EEG recording were all chosen or designed based on our pilot studies **(Supporting Figure 1)**. There were four groups: group I was sham (S), group II received a TBI and the vehicle, saline (TBI), group III received a TBI and naltrexone (T+N) and group IV did not undergo TBI and received only naltrexone (NTX). The TBI was induced by a modified Marmarou weight drop model on four-week-old male C57BL/6J mice, which is described further in the Methods section. To test whether injury to the brain after TBI alters MOR expression and whether antagonizing MOR after injury normalizes its expression, we started naltrexone treatment two hours after PTZ with additional naltrexone boluses for six days. The pro-convulsant PTZ is a GABA_A_ receptor antagonist used in various rodent models to chemically induce acute seizures. The mice were administered a sub-convulsive dose of PTZ which by itself is unable to drive the inflammatory response seen following a TBI. Rather, it is a useful test to determine seizure susceptibility in mice following the injury. Further explanation for the rational has been described previously (14–16). We first evaluated MOR expression and its status after TBI. MOR phosphorylation at specific amino acid residue sites is key for its signaling and regulation (17, 18). To elucidate the relationship between MOR phosphorylation and receptor expression, we investigated the phosphorylation status of S375, as this is one of the primary amino acid residues in MOR that is phosphorylated in response to an external stimulus.

We observed a significant increase in phospho-MOR in the neocortex (CTX) at both 8 days and 3-months post-TBI when compared to sham mice. Naltrexone treatment significantly decreased phosphor-MOR levels to near sham levels after 8 days, significantly reduced levels to sham levels by 3 months, and had no effect on its own (Fig. 1A). To further examine whether S375 phosphorylation has any effect on MOR expression, we measured MOR total protein levels. Interestingly, a very similar trend was observed in MOR protein levels to that of S375 phosphorylation. In the neocortex of TBI mice, MOR levels were significantly higher compared to sham at acute and chronic time points. Naltrexone treatment after TBI (T+N group) substantially reduced MOR protein levels. In the absence of TBI, naltrexone had no effect (Fig. 1B). The increase in MOR-positive cells shown by Western blot in the neocortex of TBI mice was confirmed by immunohistochemistry. Significantly less MOR immunostaining was observed in naltrexone-treated mice (T+N) at both time points (Fig. 1 **C-E)**. To summarize, TBI increases the expression of MORs after TBI and naltrexone reduces this effect.

**Figure 1.**
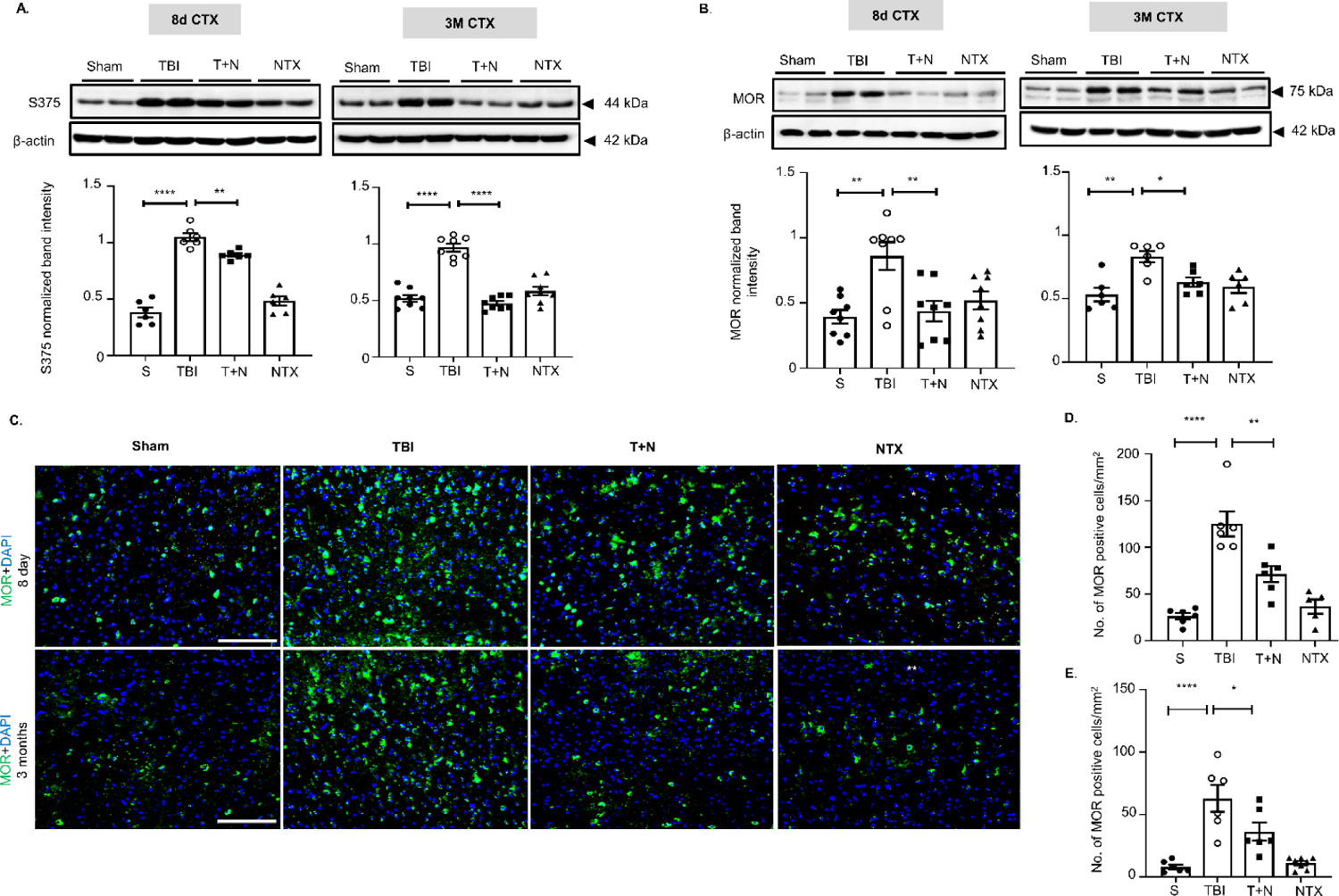
Naltrexone normalized MOR signaling in cortex following TBI. **(A, B)** Western blot analysis of phospho-MOR (S375) and MOR in the CTX tissue extract at 8 days and 3 months post-TBI. TBI increased both S375 phosphorylation (panel 1A) and MOR expression (1B) in the CTX at acute and chronic time-points compared to sham, and NTX treatment (T+N group) significantly reduced the effect of TBI. S375 after 8 days did not completely return to sham levels but by 3 months was completely back to normal levels. **(C)** MOR immunostaining in the CTX. Green represents MOR, and blue is DAPI **(D, E)**. Quantification of MOR-positive cells (with DAPI) in the CTX at **(D)** 8 days and **(E)** 3 months post-TBI, both illustrating reduction of MOR-positive cells with NTX, although not completely back to sham levels. All groups were compared to each other using one-way ANOVA with Tukey’s post-hoc test; *pC<C0.05, **pC<C0.01, \*\*\*\**p*C<C0.0001; *n* =C6–8 (3-4 sections per animal). Scale: all 100 μm. Abbreviations: S (sham); TBI (traumatic brain injury); T+N (with naltrexone treatment after TBI); naltrexone (NTX only, without TBI); neocortex (CTX).

### Naltrexone reduced TBI-induced p38 phosphorylation and nitro-oxidative stress

MOR can activate mitogen-activated protein kinase (MAPK), a pivotal regulator of inflammatory gene transcription and activation (7, 19). MAPK pathways respond to various stimuli and transduce various intra- and extracellular signals, such as stress. We hypothesized that TBI-induced stimulation of MOR triggers MAPK signaling via p38 phosphorylation and that this increased MAPK signaling activity would enhance nitro-oxidative stress and promote neuroinflammation. Immunoblots of neocortex tissue lysates detected significant upregulation of phosphorylated p38 MAPK at 8 days and 3 months post-injury.

Naltrexone, after TBI, reduced the levels of p38 MAPK phosphorylation at both time points (Fig. 2A, B). p38 MAPK phosphorylation activation correlated with phosphorylation of S375 (Fig. 1B) providing evidence of MOR activation. Two markers of nitro-oxidative stress, 3-nitrotyrosine (3-NT) and iNOS, were also substantially elevated in the neocortex of TBI mice, and elevated levels persisted for at least 3 months post-injury. Naltrexone (T+N) reduced iNOS at both time points, but 3-NT levels were only reduced at 8 days and not at 3 months (Fig. 2C, D). Thus, naltrexone has both early and partial long-term neuroprotective effects in mitigating TBI-induced nitro-oxidative stress in the neocortex.

**Figure 2.**
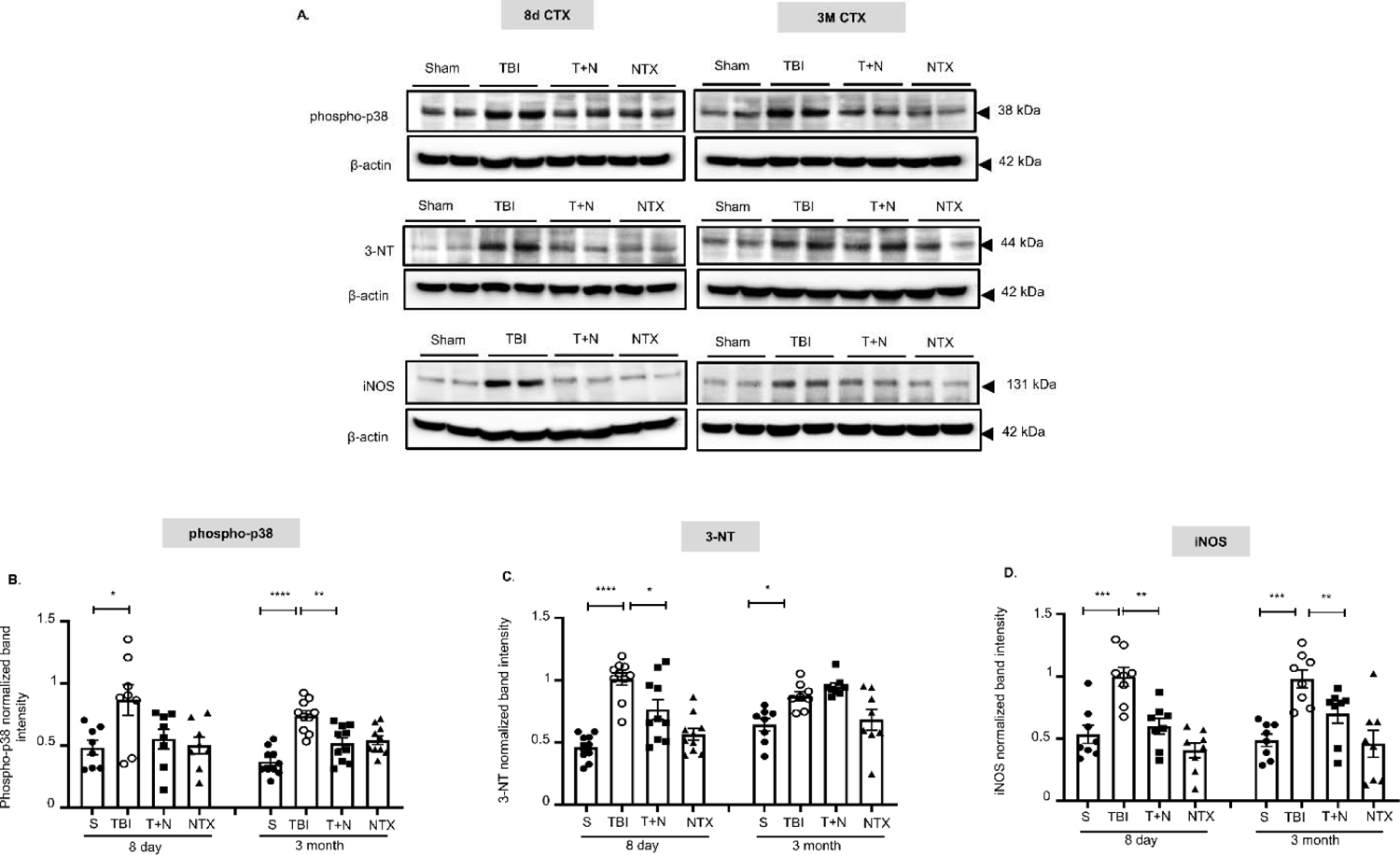
TBI-induced phospho-MAPK levels and nitro-oxidative stress markers reduced in neocortex following naltrexone administration. Western blot analysis of phospho-p38 (MAP kinase), 3-NT (a marker of protein nitrosylation), and iNOS from the CTX at 8 days and 3 months post-TBI **(A)**. Increased levels of phospho-p38, 3-NT and iNOS were detected in the CTX at both time points compared to sham **(B-D)**. NTX mitigated all biomarker levels when compared to the TBI groupC(except 3-NT at 3 months). *pC<C0.05, **pC<C0.01, ***pC<C0.001, ****pC<C0.0001; One-way ANOVA with Tukey’s post hoc test; *n*=6–8.

### Naltrexone decreased TBI-induced expression of inflammatory cytokine genes

MAP kinase phosphorylation of p38, and the subsequent stimulation and nuclear translocation of NF-kB, regulate the transcription of inflammatory genes (20, 21). Since we observed increased phosphorylation of phospho-p38 in the neocortex, we investigated whether the expression of inflammatory cytokines in the neocortex increases subsequent to the MOR-dependent phosphorylation of MAP kinase that is induced by TBI. We performed quantitative RT-PCR to determine mRNA expression of various cytokines in the neocortex.

In the TBI group compared to sham, we detected a significant increase in *IL-*1β, *TNF*α, and complementary protein *C3* transcripts and no changes in *IL-12A-B* and *IFN*γ expression at 8 days. At 3 months, mRNA expression of *IL-12A*, *IL-12B*, and *IFN*γ was significantly elevated post-TBI while expression of the other markers was normal (Fig. 3). Naltrexone treatment after TBI normalized the expression of most of the inflammatory genes evaluated, both acutely and chronically, with the exception of *TNF*α.

**Figure 3.**
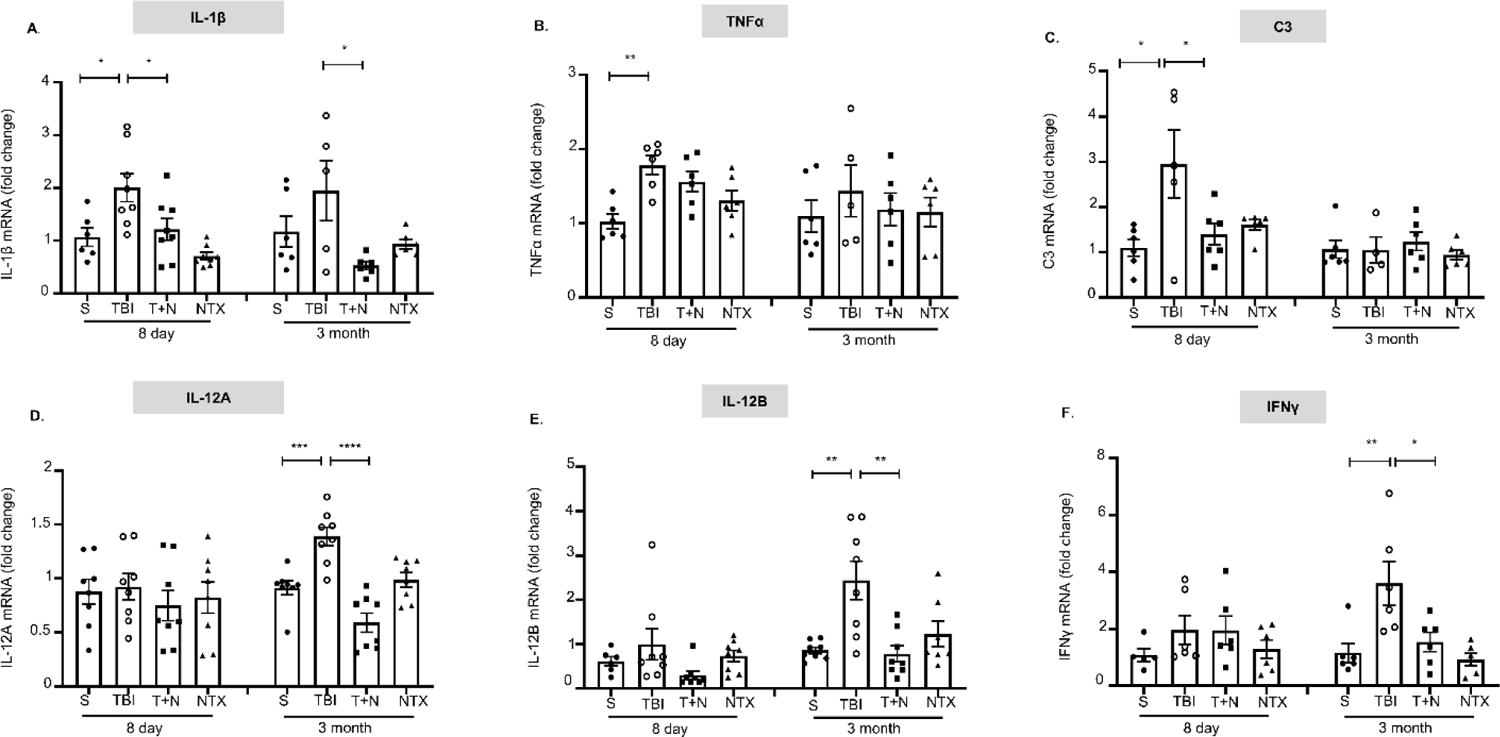
Naltrexone decreased inflammatory cytokine expression in the neocortex following TBI. mRNA levels were measured by qRT-PCR. *IL-1*β, *TNF*α, and complement protein *C3* (encode inflammatory cytokines) were significantly higher at 8 days **(A-C)**, whereas *IL-12A*, *IL-12B* and *IFN*γ mRNAs were increased at 3 months post-injury **(D-F)**. NTX treatment attenuated *IL-1*β and *C3* expression at 8 days, and decreased *IL-12A* and *IL-12B* at both time points **(*D* and *E*)**, and *IFN*γ mRNA levels **(F)** were decreased at 3 months after TBI. *pC<C0.05, **pC<C0.01; One-way ANOVA with Tukey’s post hoc test; *n*= 5-8.

### Naltrexone mitigated TBI-induced microgliosis and neurodegeneration

As the primary endogenous immune cell in the CNS, microglia are major contributors to the neuroinflammatory response following brain injury. In microglial cells, MOR stimulation promotes an inflammatory phenotype by increasing the expression of inflammatory genes and inducing neurodegeneration through oxidative stress (6, 12, 22). To assess the impact of MOR signaling on microglial reactivity and neurodegeneration, we performed immunostaining for the microglial marker, IBA1, and staining with the fluorescent dye Fluoro-Jade B (FJB). IBA1 expression correlates with microglial activation, while FJB is a well-established marker of neuronal degeneration (23, 24). Confirmation that FJB-positive degenerating cells were neurons was done by co-staining for a neuronal marker, NeuN.

Animals at 8 days and 3 months post-TBI showed acute and chronic microglial activation in brain sections. Co-labeling for IBA1 and DAPI revealed a substantial increase in IBA1-positive cells at both early and late time points post-injury, compared to sham. Naltrexone-treated animals (T+N group) had significantly fewer IBA1-positive cells in the acute and chronic phases (Fig. 4A, C). On visual inspection, it was evident that the TBI group had higher numbers of morphologically reactive microglia compared to the T+N group (**Fig 4A**). FJB staining for neurodegeneration identified higher NeuN-FJB colocalization in the TBI group at early and late time points, indicating increased neuronal degeneration. In TBI brains treated with naltrexone, there was a significant reduction in the number of FJB-positive neurons at 8 days but not at 3 months post-TBI (Fig. 4B, D) which correlates with the partial reduction in oxidative stress produced by 3-NT (Fig. 2C). Fewer NeuN-positive neurons were also detected in the TBI group neocortex, especially in the areas surrounding more densely stained FJB-positive cells, and they recovered significantly better with naltrexone treatment (Fig. 4B**; Supporting Table 3**). These results demonstrate that TBI increases microgliosis and neurodegeneration, and can be reduced markedly by blocking MOR with naltrexone, especially during the acute phase of the injury.

**Figure 4.**
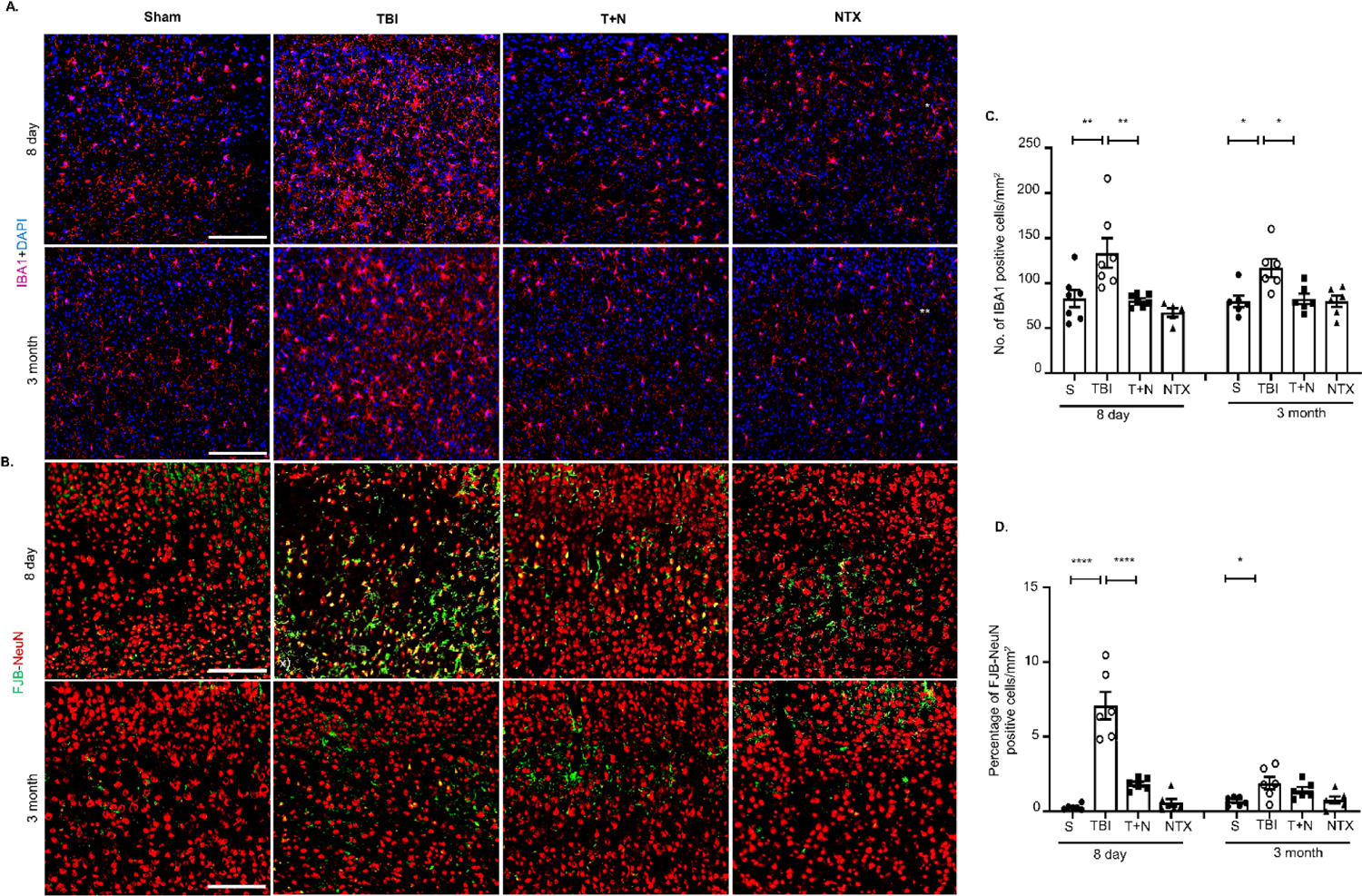
TBI-induced microgliosis and neurodegeneration in the neocortex reduced by naltrexone. **(A)** Total number of IBA1-positive cells was used to quantify microgliosis and compare the groups. IBA1 (pink) and DAPI (blue) immunostaining in panels in the CTX. Morphologically reactive microglia were primarily observed in the TBI group, while naltrexone treatment skewed microglia towards a more non-reactive phenotype as commonly observed in both sham and NTX groups. **(B)** FJB-NeuN (green-red) staining higlights NeuN-positive cells with and without FJB colocalization. Yellow cells represent neurodegeneration/stressed neurons. **(C)** Count of IBA1-positive cells and quantification of FJB-positive cells **(D)** at 8 days and 3 months after TBI. *pC<C0.05, **pC<C0.01, ****pC<C0.0001, One-way ANOVA with Tukey’s post-hoc test; *n*C=C6–8 (3 sections per animal). Scale, all 100Cμm. Abbreviations: NTX, naltrexone treatment only, without TBI; T+N, naltrexone treatment, with TBI.

### Naltrexone decreased the TBI-induced elevation of serum cytokines and chemokines

To study the systemic effects of TBI, we assessed serum cytokine levels after injury and the impact of naltrexone. Serum cytokines were quantified by multiplex ELISA at 8 days and 3 months post-TBI. In the TBI group at 8 days, cytokines IL-1α, IL-6, TNFα, IFNγ, and chemokine CCL5 were significantly elevated, whereas the anti-inflammatory cytokine IL-4 was reduced. Naltrexone, after TBI, (T+N group) effectively reduced levels of IL-1α, IL-6, TNFα, CCL5, IL-18, and IL-23 and restored IL-4 levels to normal at 8 days. At 3 months, only cytokines IL-6 and IL-12p70 were substantially elevated in the serum of injured animals.

Of the two inflammatory cytokines (IL-6, IL-12p70) in the chronic time-point, only IL-12p70 levels were significantly decreased by naltrexone (Fig. 5). As for IL-1β and IL-17 —which are the cytokines most often elevated in the serum of TBI and human epileptic patients—no significant changes were observed at 3 months nor did we detect changes in the levels of other chemokines (CCL2, CXCL5, CXCL10), growth factor VEGF, and GM-CSF or M-CSF at 3 months in all four groups (data not shown). A complete list of the cytokines evaluated for all four groups is given in **Supporting Table 2**. In summary, some inflammatory cytokines increased with TBI and decreased with naltrexone treatment post-TBI, while some anti-inflammatory cytokines decreased with TBI and increased with naltrexone treatment.

**Figure 5.**
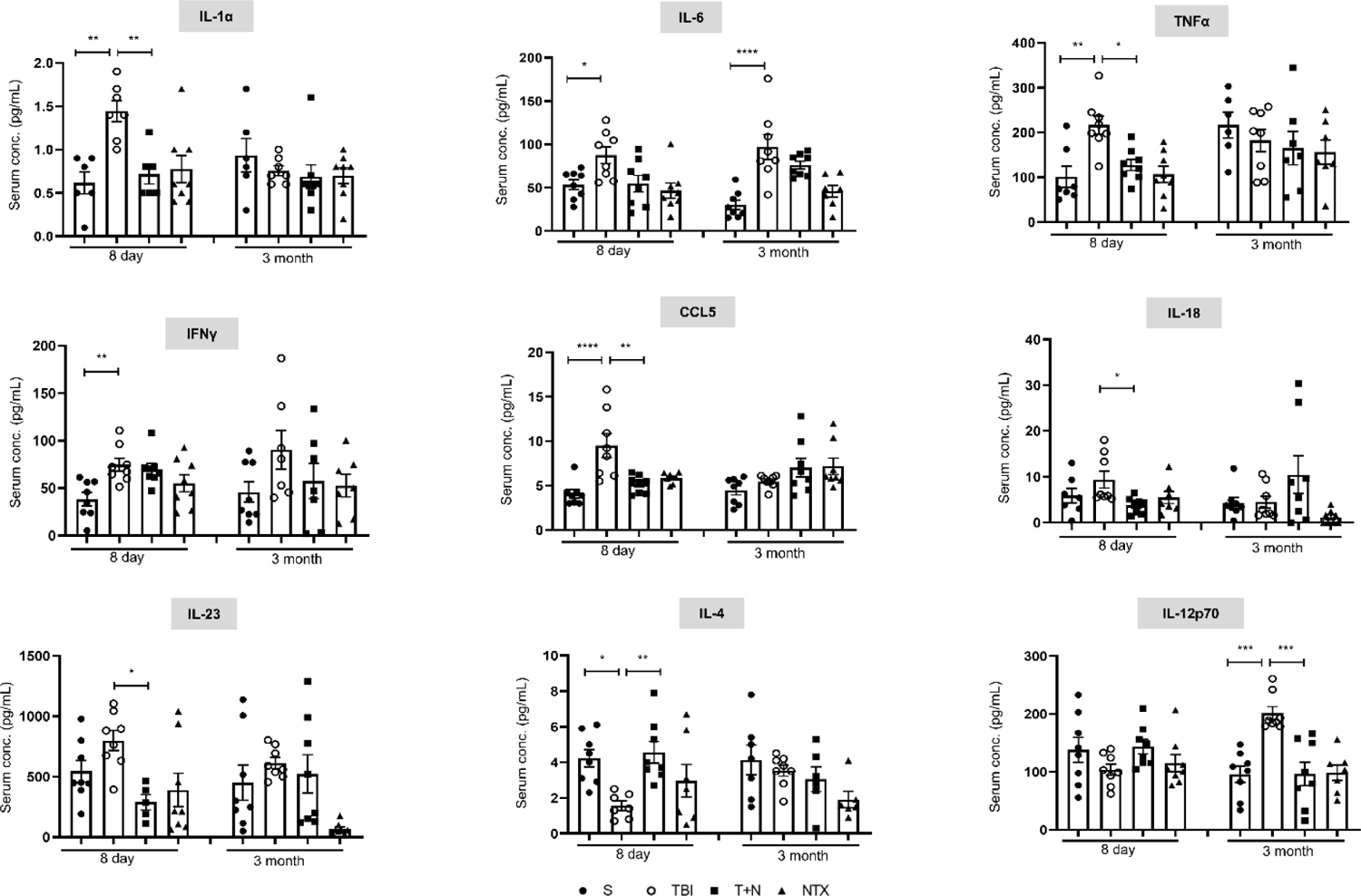
Naltrexone attenuated inflammatory and anti-inflammatory cytokines induced by TBI. The effects of NTX treatment on cytokines and chemokine levels in the serum at 8 days and 3 months post-TBI. Most of the serum cytokine levels were higher in the TBI group at the acute phase, whereas IL-6 and IL-12 were higher during the chronic phase of injury. NTX normalized the serum levels of all the inflammatory cytokines and recovered anti-inflammatory IL-4 levels. *p<0.05, **p<0.01, ****<p<0.001, ****p<0.0001; One-way ANOVA with Tukey’s post hoc test; *n*= 6-8 per group. Abbreviations: TBI, Traumatic brain injury; NTX, naltrexone.

### Naltrexone treatment of TBI brains protected the integrity of fiber tracts

Fractional anisotropy (FA) is a proxy measure of fiber tract integrity, myelination, and other factors potentially reflecting white matter pathologies in the brain (25). Decreases in FA are commonly associated with axon loss or myelination abnormalities (26). Tract-based spatial statistics analysis showed that TBI was associated with a widespread loss of fiber tract integrity. Specifically, following TBI, FA decreased significantly during the acute phase in the long interhemispheric (callosum), peristriatal, thalamic, internal capsule fiber tracts, and other regions of the brain near the injury site, indicating damage or loss of fiber tracts.

The effects on FA seen during the acute phase after TBI were also observed in the chronic phase. Strikingly, we detected no FA changes during the acute phase in the naltrexone-treated group. At 3 months, small but significant clusters of FA reduction in long interhemispheric and peristriatal fibers were observed in the naltrexone-treated group, indicating extensive protection of fiber tracts (Fig. 6 **Supporting Table 4**). Ultimately, naltrexone prevented changes in FA suggesting that naltrexone protected myelination primarily in regions such as the neocortex, peri hippocampal fiber tracts, corpus callosum, and thalamus.

**Figure 6.**
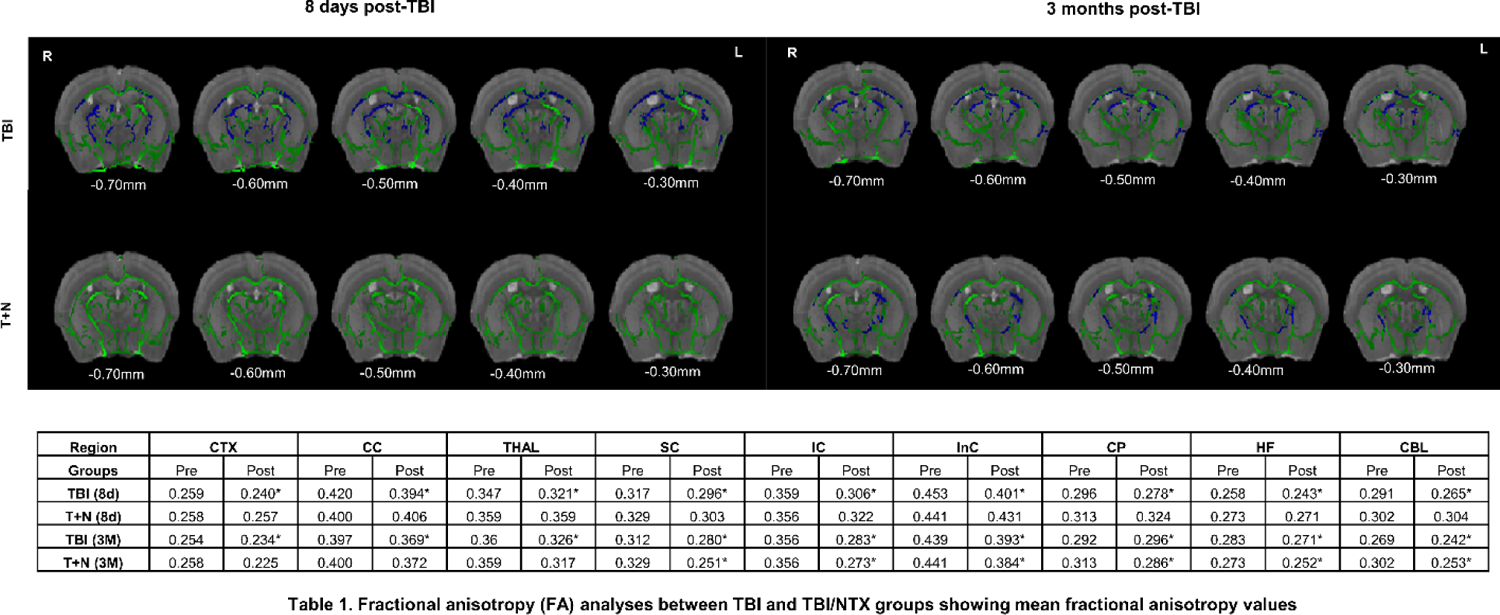
Naltrexone prevented loss of fiber tract integrity following TBI. *t-statistical* maps of whole brain diffusion tensor imaging showing differences in FA at 8 days and 3 months post-TBI. Images are overlaid onto the Waxholm template image with the mean FA skeleton shown in green. Statistics were calculated using a randomize algorithm from FMRIB software library (FSL), with 252 possible permutations and threshold-free cluster enhancement (TFCE). The images were thresholded at p < 0.05 (n=5), with significant decreases in FA shown in blue. Abbreviations: diffusion tensor imaging (DTI); fractional anisotropy (FA); tract-based spatial statistics (TBSS). **Table 1**. Mean FA values of different brain regions of the TBI and T+N groups. All the post-injury values (after TBI) were normalized against the pre-injury (before TBI) group. Therefore, the higher the value, the higher the FA, whereas lower values reflect decreased FA. The asterisk shows the difference in FA values between pre-injury (baseline scans) and post-injury scans. Abbreviations: CTX, white matter adjacent to neocortex; CC, corpus callosum; THAL, thalamic fiber tracts; SC, fiber tracts adjacent to the superior colliculus; IC, fiber tracts adjacent to the inferior colliculus; InC, internal capsule; CP, white matter adjacent to caudate and putamen; HF, white matter adjacent to hippocampus; CBL, cerebellum.

### Naltrexone decreased interictal events and prevented post-traumatic epilepsy

We used video-EEG to record the occurrence of spontaneous seizures and interictal events in mice after TBI and naltrexone treatment. Electrographic seizures were defined as high amplitude frequency discharges lasting for at least 10 seconds, with spike amplitude three times the baseline, and inter-spike interval of less than 5 seconds (27–29). The monitoring of the spontaneous seizures started at least three weeks after TBI.

The automated analysis was performed with MATLAB, and all seizures and interictal events were verified against integrated/synchronized videos associated with the EEG traces on Neuroscore software. The algorithm for automated analysis with representative EEG traces from both TBI and naltrexone-treated mice is demonstrated in our recent publication (30). Representative EEG traces showing interictal events from TBI and naltrexone-treated mice are illustrated in Fig. 7A. Twelve weeks of long-term recording revealed that naltrexone treatment significantly reduced the number of interictal events from the fourth week until the end of the observation period (Fig. 7B). In addition, none of the naltrexone-treated animals developed epilepsy, whereas 71% of the untreated animals that had TBI became epileptic (Fig. 7C).

**Figure 7.**
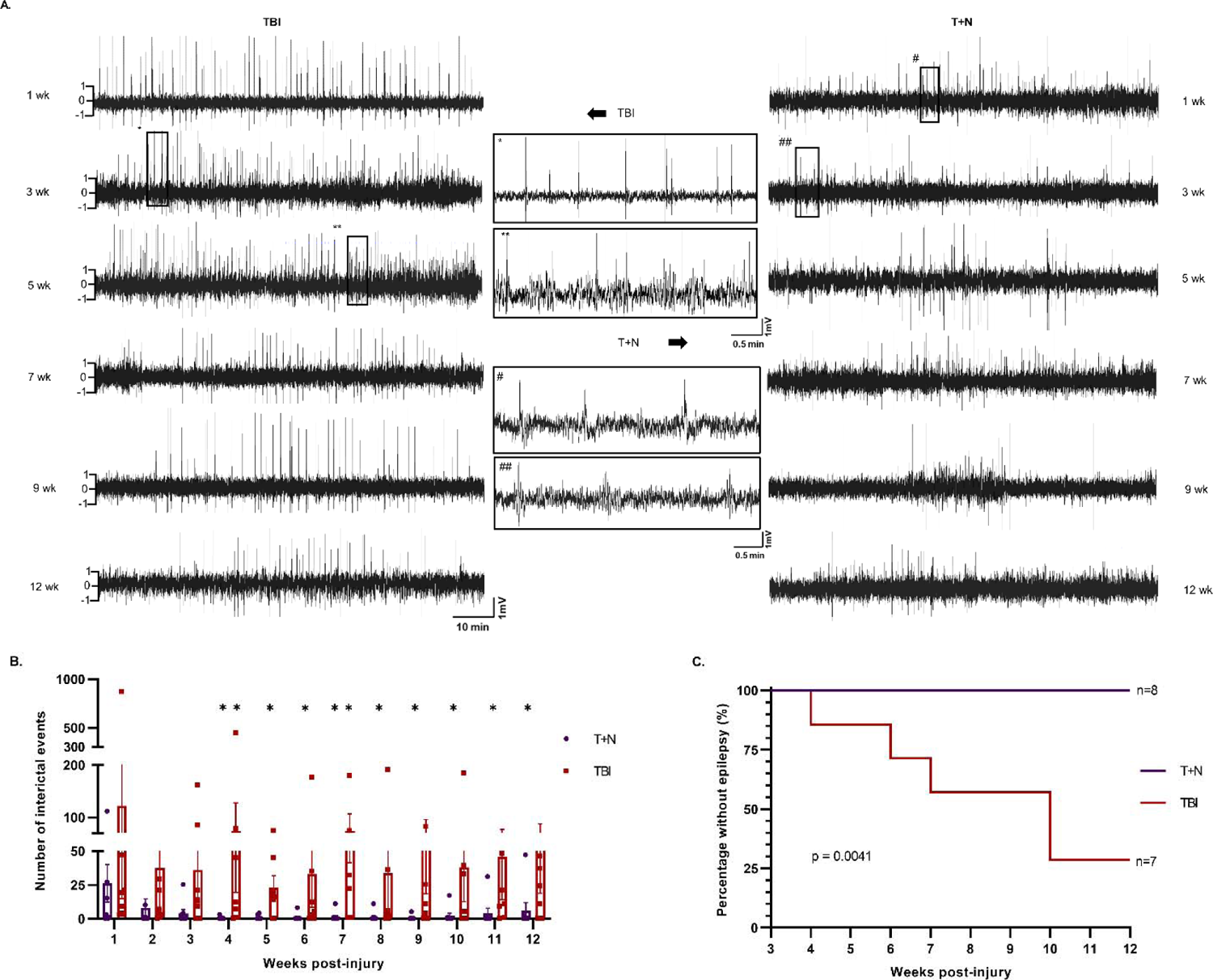
Naltrexone reduced interictal events and prevented post-traumatic epilepsy. **(A)** Segments of EEG traces showing interictal events between TBI and T+N groups. Asterisks and hashmarks represent the expanded EEG traces from the TBI and T+N groups, respectively. **(B)** The number of interictal events over 3-months was significantly reduced in the T+N group after 4 weeks and onward compared to the TBI group. **(C)** The proportion of mice with epilepsy following TBI+PTZ is significantly reduced in the T+N group compared to the TBI group. One-way ANOVA Mann-Whitney test *pC<C0.05, **pC<C0.01, *n*=7-8. Log rank P-value = 0.0041, n = 7-8 per group.

## Discussion

Naltrexone is an FDA-approved opioid antagonist widely used to manage alcohol and opioid dependence (31). Our previous studies had shown that naltrexone has anti-convulsive properties in zebrafish and mouse models, but did not address if naltrexone had anti-epileptic properties (13). To address this key translational issue, here we tested naltrexone in a mouse weight drop model of TBI with three months of video-EEG study to assay eliptogenesis, and complemented this with an analysis of a series of molecular and biochemical disease biomarkers for inflammation and inflammation-related signaling. Our results show that naltrexone strongly decreased the severity of many of the sequelae of TBI, such as neuroinflammation, nitro-oxidative stress, neurodegeneration, white matter fiber injury, interictal events and notably, epileptogenesis. With none of the mice that received NTX post-TBI developing post-traumatic epilepsy, it provides further evidence that NTX plays a role in modifying the development of epileptogenesis.

The aforementioned effects of naltrexone likely reflect selective effects on the mu opioid class of opioid receptors. Naltrexone is a potent, competitive MOR antagonist with a higher selectivity and affinity for MOR (K_i_= 1.0 nM) than for delta (DOR; K_i_= 149 nM) and kappa opioid receptors (KOR; k_i_= 3.9 nM) (32). Competitive receptor binding assays showed that the presence of a cyclopropyl methyl group on the nitrogen atom in naltrexone increases its receptor binding affinity for MOR but not KOR or DOR (33, 34). In addition to targeting Asp147 and hydrogen bond interactions at Tyr148, naltrexone also forms additional hydrophobic interactions with Lys233, causing conformational changes in MOR. These interactions cause naltrexone to bind more strongly to MOR than to other opioid receptors, making naltrexone a better and more selective antagonist for MOR (33).

Ligand binding to MOR causes a sequential and hierarchical multi-site phosphorylation of several amino acid residues leading to the receptor’s fully functional poly-phosphorylated state (35). For example, morphine stimulates phosphorylation of serine 375 (S375) and threonine 379 (T379) at lower concentrations and of T370 and T376 at higher concentratons (36). *In vivo* and *in vitro* proteomic studies on MORs revealed that application of a small amount of agonist (e.g., morphine) enhances phosphorylation of S375, a site phosphorylated early in the MOR-activation response (35). In light of this, and given that opioid receptors like MOR are known to regulate neuroinflammatory responses, we hypothesized that S375 phosphorylation increases after TBI, initiating MOR receptor activation and signaling (37). Indeed, we observed increased S375 phosphorylation in the neocortex of injured mice, which correlated well with increased overall MOR expression. Interestingly, the phosphorylation and activation of S375 and MOR persisted even through the chronic phases of injury. Antagonism by naltrexone blocked S375 phosphorylation at both acute and chronic time points, highlighting the possibility of S375 playing a role in MOR expression and signaling after TBI. Nonetheless, more research needs to be done since S375 may be among several residues controlling MOR activation.

MORs activate mitogen-activated protein kinases (MAPKs), which in turn regulate diverse cellular functions. Overactivation of MOR can cause cytotoxic oxidative stress by generating excessive reactive oxygen and nitrogen species (ROS/RNS) (6, 12, 22). Most of the damage induced by reactive nitrogen species (RNS) is triggered by iNOS and NADPH oxidase (NOX), which generate superoxide anions that combine with nitric oxide to cause neuronal death (38, 39). Early inhibition of iNOS and NOX immediately after injury prevents reactive gliosis and neuronal injury and protects the brain from oxidative damage (38, 40). In agreement with these studies, we found that nitro-oxidative stress markers were elevated in the neocortex of TBI mice and that these changes were mitigated by early treatment with naltrexone, both in the acute and chronic phases of the injury.

Oxidative stress can regulate inflammatory genes by activating MAPKs. MAPK signaling is required for the expression of iNOS and of IL-6, IL-1β, and TNFα, all inflammatory cytokines with pivotal cellular roles (41, 42). The interaction between opioid receptors and the immune system is bidirectional: receptors can be immunosuppressive or immunostimulatory. In an *in vitro* culture model enriched for primary microglia, the stimulation of MOR increased the production of IL-1β, TNFα, IL-6, and nitric oxide, causing a reactive phenotype in these cells (8). Reports have shown that chronic activation of MOR can activate microglia and astrocytes, releasing inflammatory cytokines (43). Upregulation of these cytokines in glial and immune cells has also been linked to brain neurodegeneration in various chemo-convulsant models of temporal lobe epilepsy (44–46). Likewise, in our experimentally injured mice, IL-1β, TNFα, and other cytokines were upregulated in the neocortex at early and late time points post-injury and, concurrently, their brains developed gliosis and neurodegeneration. At the same time, MOR protein and S375 phosphorylation levels also increased. Strikingly, inhibiting MOR with naltrexone reduced inflammatory cytokines in the brain and serum and prevented neuroinflammation, suggesting that blocking MOR with naltrexone decreases immune responses and inflammation.

TBI changes the anterior white matter structure broadly, which can be visualized through fractional anisotropy (FA)—a measure of anisotropic water diffusivity in the brain that provides a read-out of fiber organization, orientation, and the degree of myelination (47). In various closed-head injury models, numerous neuroimaging studies reported widespread damage to the white-matter structures and discontinuity in fiber reconstructions in different brain regions after injury (48–50). Studies on epilepsy models have also reported a decrease in FA in major bundle fibers, in the hemisphere ipsilateral to the seizure origin and in the corpus callosum, and in white matter adjacent to the neocortex and hippocampus (47, 51, 52). Since trajectories of FA changes are likely predictive of outcomes after TBI, we used FA to evaluate changes in the whole brain after TBI and with naltrexone therapy. Consistent with the previous studies, we found reductions in FA after TBI, especially in the white matter adjacent to the neocortex, corpus callosum, and internal fiber tracts, both during the acute and chronic phases (53). Naltrexone treatment prevented the changes in FA in most regions, especially in the regions proximate to the neocortex, peri hippocampal fiber tracts, corpus callosum, and thalamus, which suggests that naltrexone protects myelination. The mechanisms by which naltrexone protects myelination and maintains white-matter fiber integrity in the brain are unknown. Persistent brain inflammation causes tissue damage and white-matter degeneration. Perhaps by attenuating inflammation, naltrexone protects white matter from deterioration. Yet, we should also consider other factors, such as changes in diffusivity, cerebral blood flow autoregulation, blood-brain-barrier disruption, and genetic modulators. Additional studies are required to confirm the correlations of these factors with white-matter damage after the injury and to explore mechanisms underlying naltrexone’s neuroprotective effects.

Although the persistent occurrence of interictal spiking does not always lead to epilepsy, its occurrence over a long period of time can serve as a crucial diagnostic biomarker for epilepsy (54, 55). In our study, we observed interictal events over the twelve-week observation period after TBI. Notably, these discharges were reduced by naltrexone, and epilepsy did not develop for up to 3 months in the naltrexone-treated group. While Naltrexone’s target, MOR, plays a known but incompletely understood protective role in epilepsy, this receptor subtype has also been implicated in the pro-convulsive actions of morphine (56, 57). Our observations provide evidence that MOR may function in the pro-convulsive capacity after TBI and that naltrexone can block this effect.

The increased spiking observed in the TBI group in our model could be linked, in principle, to either enhanced neuronal excitability or decreased inhibition. As MORs are widely expressed on GABAergic interneurons, they can reduce neuronal GABAergic activity, causing disinhibition (58–60). This may facilitate excitatory signaling, thereby enhancing seizure susceptibility. Another plausible explanation is that MOR increases excitability in pyramidal cells by closing K+ channels and increasing NMDA receptor- and L-type Ca^2+^ channel-mediated Ca^2+^ entry (61, 62). In our prior publication studying naltrexone’s anti-convulsant effect in mouse brain slices and in a zebrafish model, seizure-like events were induced by blocking GABAergic activity with pentylenetetrazol (13). Thus, our prior results suggest that naltrexone may act by decreasing excitability rather than altering inhibition.

Collectively, our previous and our current study indicate that naltrexone has anti-convulsive, anti-epileptic and anti-inflammatory properties. A limitation of our study is that it does not answer whether naltrexone’s neuroprotective effect is due to decreased inflammation, by directly altering neuronal excitability, or by a combination of both. Future studies will be designed to address this important issue. Another limitation of our study is the duration over which post-traumatic seizures were evaluated. Future studies will involve longer post-TBI periods to assess better the long-term effect of naltrexone, especially its anti-eliptogenic effect.

In conclusion, our findings illustrate the neuroprotective effects of naltrexone in a mouse model of post-traumatic epilepsy when the drug was administered starting one day after the brain injury. Mechanistically, naltrexone has strong anti-inflammatory properties and reduces neuroinflammation by targeting reactive gliosis, oxidative stress (ROS/RNS), inflammatory cytokine production, neurodegeneration, and epileptiform discharges and, consequently, modifying the development of post-traumatic epilepsy, either slowing or preventing eliptogenesis.

## Methods

### Experimental groups, induction of traumatic brain injury, and drug treatment

Mice (n=95) were randomly divided into four groups. Group I, sham (S), received no treatment; Groups II (TBI) and III sustained a TBI, after which Group III received naltrexone (T+N); and Group IV received naltrexone, without TBI (NTX group). To induce TBI, a Marmarou WD model was employed on four-week-old male C57BL/6J mice, with slight modifications to minimize variability (63, 64). Animals were habituated for 2 h prior to injury and anesthetized using gaseous isoflurane. The depth of anesthesia was assessed by a toe pinch test. An eye ointment was applied, and 1 mg/kg meloxicam was administered subcutaneously prior to WD. A free fall weight of 50 g was dropped from the height of 45 cm so that the weight would impact the midline of the skull between the ears. The weight and height of the impact were standardized based on our pilot studies, taking mortality and injury severity into account **(Supporting Figures 1 and 2**). Post-impact, animals were immediately placed on a heating pad, and the time taken by each mouse to regain consciousness was recorded. One hour after injury, the neurobehavioral status of the mice was assessed by a Neurological Severity Score (NSS) test **(Supporting** Figure 2). At the end of the assessment, softgel, Nutrical and 0.5 mL saline was given to all the animals. For non-telemetry animals (without transmitter), a second treatment, referred to as the PTZ test, was performed on day 2 post-TBI, and for telemetry animals (with transmitter), the same PTZ test was performed on day 5 post-TBI. The choices of dosage, route, and treatment regimen for the PTZ test were decided based on our pilot data, the toxicological and drug safety profile studies of PTZ, and the drug’s pharmacological properties (65–71). This PTZ test consisted of a sub convulsive dose of PTZ, given as a second “hit” after the initial weight drop at the times specified above for the non-telemetry and telemetry groups. Since this dose has no convulsive action, the weight drop is the primary experimental insult. The administration of sub-convulsive doses of chemoconvulsants is a well-established and frequently used method to determine seizure susceptibility in mice and rats following TBI (14–16).

Naltrexone treatment (40 mg/kg, subcutaneous) was initiated two hours after PTZ injection, and was continued every 12 hours for three days and then once daily for four days (Fig. 8). EEG was recorded continuously for 8 days after TBI, and then intermittently (8 h/day) for up to 3 months. Animals were euthanized at 8 d and 3 months post-TBI. PTZ and naltrexone were freshly prepared in saline at a 5 mg/mL concentration. All experiments on animals were performed under the Institutional Animal Care and Use Committee guidelines, University of Iowa, USA.

**Figure 8.**
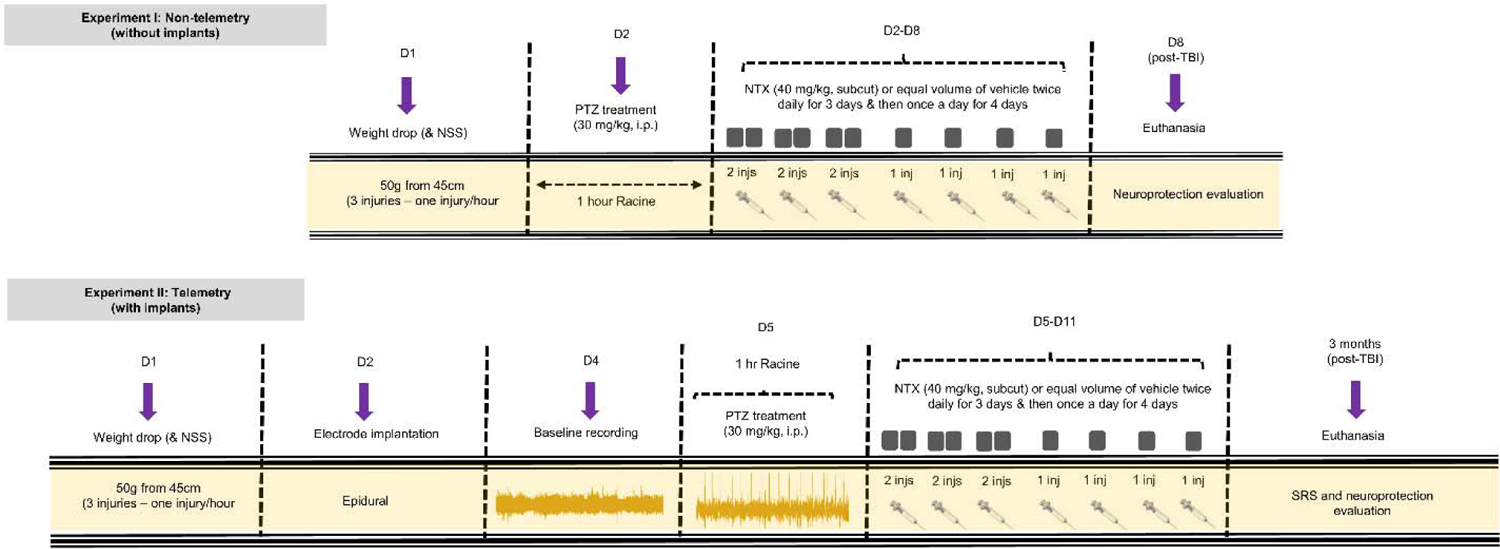
Experimental protocol to model TBI and assess naltrexone therapy. Experimental design illustrating the two experimental groups, which are designated the non-telemetry (experiment I, without implants) and telemetry (experiment II, with implants) groups, and detailing their respective treatment regimens and endpoints. For non-telemetry animals, traumatic brain injury was induced on day 1 (D1) and the PTZ-test was performed on day 2, whereas for telemetry animals, the PTZ-test was performed on day 5 post-TBI, i.e., after the electrode implantation (day 2). Naltrexone treatment was initiated 2 hours after administration of the sub-convulsive dose of PTZ. The monitoring of the spontaneous seizures (experiment II) started three weeks after TBI. Animals were euthanized at 8 d (for experiment I) and 3 month (for experiment II) post-TBI.

### Electrode implantation, radiotelemetry setup and video-EEG recording

The detailed procedure for electrode implantation, radiotelemetry setup and recording are described in our previous and recent publications (28, 30, 72, 73). Mice were placed on a heating pad and anesthetized using gaseous isoflurane. The head was shaved, artificial tear ointment was applied (Aventix, CAT: 13585) and meloxicam (1 mg/kg, subcutaneously) was administered prior to surgery. A midline incision was made from the mid-dorsal region of the head toward the neck to expose the skull. A subcutaneous pocket was created in the flanking region for the HD-X02 PhysioTel^TM^ radio-transmitter (Data Science International, CAT: 270-0172-001). Bilateral burr holes were drilled into each hemisphere of the brain and electrodes were placed epidurally in a V-shaped pattern. The electrodes were secured in place using dental cement. Surgical clips and sutures were used to close the incision. After surgery, antibiotic Baytril (5 mg/kg, subcutaneous) was given to prevent bacterial infection. Topical triple antibiotic ointment was applied on the incision site, and 1 ml saline was given to replenish lost electrolytes. Mice were then placed in an empty cage on a heated pad until they regained consciousness and became ambulatory.

One day after electrode implantation, mice were moved into the telemetry room and 24 hours of baseline EEG was recorded using a PhysioTel^TM^ radiotelemetry device. Animals were individually caged and placed on PhysiolTel^™^ receiver model RPC-1 receiving pads. Pads were spaced out on tables with aluminum shields placed between pads in order to eliminate signal crossing between pads. These receiving pads were connected to a Matrix 2.0 for PhysiolTel^™^ for data exchange with the PC through Ponemah, version 6.51. Four AXIS M1145-L Network Cam Kits and Media Recorder 4 software, version 4.0, were used to capture video to help create eight video-synchronized EEG traces. One camera kit was able to record two receiver pads at a time. The Ponemah software combines all data from the Media Recorder software and the Matrix 2.0 to create video-synchronized, two channel, EEG traces with the sampling frequency of 1000 Hz with a filter cutoff of 100 Hz and full scale at 10 mV. Videos were captured at 20 frames/second with a resolution of 1280×720. Files were saved directly onto a 4TB Seagate hard drive.

Following the acclimation period and baseline recording, the PTZ test was performed by administering a sub-convulsive dose of 30 mg/kg intraperitoneally. Following one hour of behavioral observation (or 2 h post-PTZ dose), naltrexone (40 mg/kg, subcutaneous) or vehicle treatment was initiated and was given twice a day for three days and then once a day for four days. For the initial phase of EEG recording (the first seven days following TBI by Weight Drop), video-synchronized EEG was recorded for 24 hours/day, and then, in the subsequent phase, EEG was recorded intermittently for up to three months. For this period, EEG was recorded for 8-10 hours a day for five days a week, followed by video recordings overnight and over the weekends. All the seizures were verified against the power spectrum (high-frequency gamma bands), electromyogram, and the videos synchronized with EEG.

### Quantification of electrographic spikes and seizures

We developed a novel MATLAB-based algorithm that analyzes spiking activity and interictal events to quantify mouse EEG data. Mouse EEG data was imported into MATLAB. Baseline EEG activity was determined autonomously for each individual data file. The lower threshold for spike detection was specified as twice the baseline activity (74). The upper threshold for spike detection was set at 1500 μV, as spikes larger than 1500 μV often arise from electrical interference (27, 28). Next, EEG spikes were identified by the algorithm and were passed through a set of filters to remove any spike that failed to meet the spiking criteria (minimum peak height of 200 µV; maximum peak width of 200 ms). The remaining algorithm code quantified interictal events by analyzing clusters of spiking activity. These events were again passed through the filters, removing any event that failed to meet the interictal event criteria (28, 75). Each spike in an event has amplitude 3X times the baseline, with >3 spikes in an event and an inter-spike interval of 10 s (75, 76). Electrographic seizures were also identified through a similar pipeline, that is, where lower detection threshold was twice the baseline, upper detection ≤ 1,500 μV and an inter-spike interval of ≥ 100ms. Events with increased duration and magnitude were manually reviewed using Neuroscore software. The detailed procedure for our algorithm-based automatic quantification of electrographic spikes and seizures has been published recently (30).

### Immunohistochemistry, microscopic imaging, and cell quantification

Animals were transcardially perfused initially with 1XPBS for five minutes, followed by 4% paraformaldehyde (PFA) at 8 days and 3 months post-TBI. Blood was collected, stored at 21°C for 10 minutes, spun at 2,000xg at 4°C, and serum was collected. After fixation, brains were dissected out and post-fixed in 4% PFA overnight at 4°C. The following day, fixed brains were cryopreserved in 30% sucrose for 3-4 days at 4°C or until they settled to the bottom of the vial. All tissues were then embedded in gelatin type B, wrapped in saran wrap and stored overnight at 4°C. The next day, brain blocks were prepared by snap-freezing the gelatin-embedded tissue blocks in liquid nitrogen, using 2-methylbutane, and stored at −80°C. Brain blocks were serially sectioned into 12 µm thickness sections using a CryoStar NX70 cryostat with 3-4 sections/slide mounted sequentially at approximately 375 µm apart, representing rostral to caudal parts of the brain. The section sampling and collection method has been described in detail by Puttachary et al. (2016). For immunohistochemistry (IHC), sections were washed with 1XPBS for 45 minutes and then blocked with 10% donkey serum containing 0.2% Triton X-100. After blocking for an hour at room temperature (RT), brain sections were stained with primary antibodies of interest and were incubated overnight at 4°C. The following day, after washing with 1XPBS, sections were immunostained with the appropriate secondary antibody (FITC- or CY3-conjugated) for an hour at RT, washed with 1XPBS again and then were mounted with vectashield 4’,6-diamidino-2-pheny-lindole (DAPI) for nuclear staining (72, 73, 76). Descriptions and dilutions of the primary and secondary antibodies are listed in Supporting Table 1. To determine the extent of neurodegeneration, we performed Fluorojade B staining (FJB), co-immunostained with NeuN. For FJB staining, sections were incubated in 0.006% potassium permanganate solution for 10 mins. After washing three times with distilled water (30 seconds for each wash), slides were submerged in 0.001% FJB+0.1% acetic acid solution for 10 mins in the dark. After washing, slides were air-dried in the dark at room temperature, cleared with xylene and mounted with Surgipath Acrytol (Surgipath, Leica Biosystems, IL) (72, 73, 76). To eliminate variability, brain sections from all the groups were prepared and stained simultaneously.

Images were taken using a Hamamatsu C13440 Digital Camera, ORCA-Flash 4.0 and Pika Allies vision, on an Olympus virtual slide scanner with a 4X DAPI overview scan. FITC and TRITC exposures were 101.45 ms and DAPI exposure was 70 ms, and the images of all the regions of interest (ROI) were taken at 20X. For magnification, multiple focus points were placed within the ROIs to attain clear pictures by adjusting the clarity points manually. After the slides were scanned, images were converted to TIFF files for quantification using ImageJ. All the images were counted bilaterally and manually from 3-4 sections/animal at bregma levels −1.45, −1.95, and −2.35 at 20X magnification using ImageJ from a known area. IHC data was expressed as positive cells/mm^2^ and the outcome determined by averaging the total number of cells among the sections.

### Western blotting

The cortical tissues were dissected from the mice at 8 days and 3 months post-TBI and snap-frozen in liquid nitrogen. Tissues were homogenized and lysed in NET100 buffer containing 5M NaCl_2_, 0.5M EDTA and 1M Tris pH 8 with 1% protease and phosphatase inhibitor (ThermoFischer Scientific, USA). The supernatant was collected and stored at −80°C for later analysis. The protein concentrations from tissue lysates were determined using the Bradford assay kit (Biorad, USA; Cat# 5000006). Equal amounts of protein (30 μg) were loaded in the wells of 7.5% −10% precast denaturing polyacrylamide gels along with a molecular weight marker. The gels were run at RT at 80 V for 1 h, and then at 120 V until the bromophenol dye reached at least 0.5 cm from the bottom of the plate. Protein transfer to the nitrocellulose membrane was carried out by rapid transfer: the transfer sandwich containing the gel and membrane was placed into a Trans-Blot Turbo Transfer System (Biorad, USA), which was then run at 25 V for 9 minutes. Next, the membrane was washed with 1XTris-Buffered Saline (TBS) for 10 minutes and transferred to 5% nonfat dry milk in Tris-Buffered Saline with Tween 20 (1XTBST) (Sigma; cat# T-9039) for blocking for 1 hour at RT. After the blocking step, the blots were incubated with primary antibodies of interest overnight at 4°C. Descriptions and dilutions of the primary antibodies are listed in Supporting Table 1. The following day, the membranes were washed with TBST and incubated with Peroxidase AffiniPure anti-rabbit and mouse IgG antibodies (1:10,000, Jackson ImmunoResearch, USA) for 1 h at RT followed by further washes with TBST as described earlier. 5% milk and/or 5% BSA in TBST was used as a diluent for both primary and secondary antibodies. Protein bands were detected using SuperSignal^TM^ West Pico Plus chemiluminescent substrate and detected using a MyECL imager. All bands were normalized against β-actin and quantified using densitometric analysis on Image J.

### Quantitative real time-polymerase chain reaction

Gene expression of inflammatory cytokines was determined by quantitative real-time PCR. RNA was extracted using the trizol-chloroform method per manufacturer’s instructions (ThermoFisher Scientific). 1 µg of RNA was used for cDNA synthesis using SuperScript® III First-Strand Synthesis kit, yielding high-quality single-stranded cDNA. cDNA synthesis reactions were performed as follows: primer annealing at 70°C, 10 min; hold at 15°C for 5 mins; then reverse transcription at 42°C, 50 min; 70°C for 15 min and hold at 15°C. The following day, plate reactions were prepared using amplified cDNA diluted in ultrapure water (1:15) with a 2X universal master mix. Plates were run for quantitative real-time PCR (qRT-PCR) using pre-validated qPCR primers for the genes of interest **(Supporting Table 1**) on a QuantStudio^TM^ 7 Pro using FAM-MGB_VIC-Tamra fluorescent labels, as follows: 10 min, 95°C; then 95°C for 15 s, 60°C for 1 min (40 cycles). GAPDH was used to normalize all the genes. The fold change in mRNA expression was determined using cycle threshold (Ct) values, and results were expressed as fold difference from controls.

### Multiplex cytokine assay

A multiplex assay using a Quantibody Custom Mouse Cytokine Array (QCMCA) was performed by RayBiotech, on the serum samples from 8 day and 3-month post-TBI mice (CAT: QAA-CUST, RayBiotech Inc, GA, USA). The core QCMCA technology uses glass slides onto which protein targets are attached in an array format, allowing for multiple detection of 32 (or more) analytes in the same sample at a given time. Briefly, cytokine standard dilutions were prepared from a lyophilized cytokine standard mix after reconstituting with 500 μl sample diluent (standard 1). 200 µL of sample diluent was added to six different microcentrifuge tubes (standards 2-7). Serial dilutions were performed by adding 100 μl of standard 1 to standard 2, and then from standard 2 to tube 3, and so on. For blocking, 100 μl of sample diluent was added to each well and incubated for 30 minutes. After the buffer was decanted, 100μl of standard cytokines or samples were added and incubated at RT for 1-2 hours. Post-incubation, samples were washed five times for five minutes each with 150 μl of 1X wash buffer. Detection antibody was reconstituted with 1.4 mL of sample diluent, and 80 μl of detection antibody cocktail was added to each well and incubated at RT for 1-2 hours. After incubation, samples were decanted, and each well was washed with 150 μL of 1X Wash Buffer I and twice with 150 μL of 1X Wash Buffer II. After washing, 80 μL of dye-conjugated streptavidin was added to each well, covered with aluminum foil, and incubated for an hour at RT. Wells were washed five times using wash/drying/wash cycles with 1X Wash Buffer I and II. Signals were visualized with a laser scanner equipped with Cy3 wavelength channel such as Axon GenePix or Innopsys Innoscan. Data were extracted using GAL files and converted to Excel files.

### Acquisition and analysis of diffusion-weighed MRI Image acquisition

Mice from both groups, TBI and T+N, were scanned before TBI, and at 8 days and 3 months post-TBI. Imaging was performed on a GE/Agilent Discovery 901 7-Telsa pre-clinical scanner. Mice were anesthetized with isoflurane (3% induction, 1.5% maintenance) in oxygen, and depth of sedation was monitored via the respiratory rate with an MR-compatible monitoring system (SA Instruments, Inc., New York). MRI imaging acquisitions included a high-resolution 3D anatomical scan to derive structural information and a 32-direction diffusion tensor imaging (DTI) scan to investigate microstructural integrity. The 3D anatomical scan was acquired with three-dimensional fast imaging employing a steady-state acquisition pulse sequence. A 224 × 192 × 112 matrix was acquired over a 25 mm × 25 mm × 20 mm field of view resulting in a native voxel resolution of 0.11 mm × 0.13 mm × 0.18 mm at a TR/TE = 5.8ms/2.8ms, flip angle 30°, and 3 signal averages. The diffusion-weighed scan utilized an 8-shot 2D segmented echo-planar imaging (EPI) to a 128 × 128 matrix over a 25 mm × 25 mm field of view with 20 slices at 0.8 mm thickness. The scan acquired 15 diffusion directions with b = 1000 s/mm² along with 2 T2 (b = 0) images with TR/TE = 4000ms/18.4ms.

### Image preprocessing and TBSS analysis

After acquisition, diffusion-weighed images were converted from DICOM to NIFTI format using DCM2NIIX and examined for quality. Bias field correction was applied using N4BiasFieldCorrection from Advanced Normalization Tools (ANTs), and images were resampled into a voxel resolution of 0.2 mm^3^ for use in the processing pipeline (77). We performed voxel-wise statistical analysis of collected fractional anisotropy (FA) data using a version of Tract-Based Spatial Statistics (TBSS), from FSL developed in house and optimized for mouse imaging (77, 78). Using the diffusion toolbox (FDT), a tensor model was fit to the diffusion data to generate FA images, which were brain-extracted using hand-drawn masks. FA data for all animals were then aligned into Waxholm space using the nonlinear registration tool FNIRT from FSL (79). Next, mean FA images were created and thresholded at a value of 0.2 to delineate major fiber tracts and create a mean FA skeleton representative of the centers of all fiber tracts in the data. Each subject’s aligned FA data was projected onto the mean FA skeleton, and the resulting data was used to perform voxel-wise cross-subject statistics using the randomize tool from FSL, with 252 permutations and threshold-free cluster enhancement (TFCE) at *p*<0.05 to check for significance in the contrasts.

### Methodological rigor

The experimenters were blind to the experimental groups until after the data analyses were completed. We followed pre-determined criteria to exclude animals from the data analyses. The criteria set were: i) if animals did not respond to the predetermined injury (increased loss of consciousness [LOC] and neurological severity score [NSS] compared to sham); ii) if animals died during the course of the experiment; and, iii) if animals did not regain bodyweight within 8– 10 days and if EEG electrodes were displaced during the recording period. We took measures to minimize variables by: i) randomizing the animals based on a predetermined weight (≥12 g mice) and age range (4 weeks) before the start of the experiment; ii) quantifying injury severity during the weight drop by both direct observation and offline video analysis and using at least two independent observers; iii) acquiring ∼24 h of baseline EEG data, covering day-night cycles, to normalize post-TBI EEG from the same animal; iv) where appropriate, implementing the first two of the three principles of reduction, refinement, and replacement (3Rs) by adopting a refined, predetermined method of TBI induction, which reduced the mortality rate and minimized variability in TBI severity between animals and groups; and, v) determining the optimum concentration of the primary antibodies by serial dilution and validating their specificity using neutralizing antibodies appropriate to the primary antibodies.

## Statistics

Normality of the data was tested using the Shapiro-Wilk and Kolmogorov-Smirnov tests. Unpaired *t*-tests were used for parametric comparisons, while the Mann-Whitney U was used for unpaired non-parametric comparison. One-way repeated analysis of variance (ANOVA) was used for multiple comparisons of parametric data with Tukey’s post-hoc analysis. Epilepsy incidence analysis was performed utilizing Kaplan-Meier analysis with log rank test. Statistical significance was set to *p <* 0.05. Prism 9 was used for data analysis.

## Data Availability

Data will be available upon request. Supportive analytic code is available here https://github.com/Jackson-Kyle-CCOM/Automated-EEG-Algorithm.

## Study Approval

Animal procedures were performed in accordance with NIH guidelines and approval from IACUC of the University of Iowa.

## Author contributions

SS and AGB contributed to the conception and design of the study. SR, SS, GT, ZP, KJ, DT, AW, DK contributed to data acquisition, and all authors contributed to the analysis. SR, SS, and GT contributed to drafting the manuscript and preparing the figures. All authors contributed to manuscript editing. AGB was responsible for project supervision, administration, and funding acquisition. The order of co-first authors was determined by final contributions to the writing and experiments.

## Acknowledgments

The work was supported by NIH 5R01NS098590, 1P50HD103556-01A1 and the UIDM/Tross Family grant to Dr. Alexander G. Bassuk, by NIH 5R01NS115800 and an Iowa Neuroscience Institute grant to Dr. Joseph Glykys, by NIH K08NS110829 and an Iowa Neuroscience Institute grant to Dr. Elizabeth Newell, and by NIH R01EY031952, R01EY030151, R01EY024665 and R01EY025225 grants to Dr. Vineet B. Mahajan. The authors would like to thank Noah Gilkes for assisting with qRT-PCR. The authors would also like to thank Michael Rebagliati of the Scientific Editing and Research Communication Core at the University of Iowa Carver College of Medicine for editing the manuscript. T.N.-J. was supported by the Andrew H. Woods Professorship.

## Notes

https://github.com/Jackson-Kyle-CCOM/Automated-EEG-Algorithm

### Competing Interest Statement

The authors have declared no competing interest.

