## Supplemental Material for "Neuroprotective Effects of Naltrexone in a Mouse Model of Post-Traumatic Epilepsy"

**Table of contents**

**Supporting Tables**

Supporting Table 1	Reagents‘ Table

Supporting Table 2	Serum multiplex cytokine assay table

Supporting Table 3	Quantification table for NeuN positive cells

Supporting Table 4	Fractional anisotropy analysis

**Supporting Figures**

Supporting Figure 1	Standardization of surgical and drug administration protocol

after TBI

Supporting Figure 2	Neurobehavioral status of mice after TBI

**Raw Western Blots**

Western Blot 1	S375

Western Blot 2	MOR

Western Blot 3	Phospho-p38

Western Blot 4	3-NT

Western Blot 5	iNOS

**Supporting Table 1.** Reagents

| **Antibody** | **Species/Clone** | **Used For** | **Dilution** | **Catalog #** | **Vendor** |
| --- | --- | --- | --- | --- | --- |
| Anti-MOR | Rabbit polyclonal | IHC | 1:1500 | AB1580-I | Millipore Sigma |
| Anti-IBA1 | Goat polyclonal | IHC | 1:400 | NB100-1028 | Novus Biologicals |
| Anti-NeuN | Rabbit polyclonal | IHC | 1:400 | ab104225 | Abcam |
| Anti-MOR | Rabbit polyclonal | WB | 1:1000 | AB1580-I | Millipore Sigma |
| Anti-Phosphor-p38 MAPK | Rabbit monoclonal | WB | 1:1000 | 4511S | Cell Signaling |
| Anti-3-NT | Mouse monoclonal | WB | 1:1000 | ab61392 | Abcam |
| Anti-iNOS | Rabbit polyclonal | WB | 1:800 | NB300-605 | Novus Biologicals |
| Anti-β-actin | Mouse monoclonal | WB | 1:5000 | A2228 | Sigma-Aldrich |
| FITC-conjugated | Anti-goat | IHC | 1:50 | 705-095-147 | Jackson Immunoresearch |
| FITC-conjugated | Anti-Mouse | IHC | 1:80 | 715-095-150 | Jackson Immunoresearch |
| FITC-conjugated | Anti-rabbit | IHC | 1:100 | 711-095-152 | Jackson Immunoresearch |
| CY3-conjugated | Anti-rabbit | IHC | 1:200 | 711-165-152 | Jackson Immunoresearch |
| CY3-conjugated | Anti-goat | IHC | 1:200 | 705-165-147 | Jackson Immunoresearch |
| Peroxidase AffiniPure Goat Anti-rabbit | Anti-rabbit | WB | 1:10000 | 111-035-144 | Jackson ImmunoResearch |
| Peroxidase AffiniPure Goat Anti-mouse | Anti-mouse | WB | 1:10000 | 115-035-003 | Jackson ImmunoResearch |

| **Primer** | **Used For** | **Catalog #** | **Vendor** |
| --- | --- | --- | --- |
| IL-1β | qPCR | Mm00434228_m1 | ThermoFisher Scientific |
| TNF-α | qPCR | Mm443258_m1 | ThermoFisher Scientific |
| IFNγ | qPCR | Mm01168134_m1 | ThermoFisher Scientific |
| C3 | qPCR | Mm01232779_m1 | ThermoFisher Scientific |
| IL-12A | qPCR | Mm00414165_m1 | ThermoFisher Scientific |
| IL-12B | qPCR | Mm00434174_m1 | ThermoFisher Scientific |

Primary and secondary antibodies used for immunohistochemistry and Western blotting. Gene primers used for Quantitative real-time PCR.

**Supporting Table 2.** Multiplex cytokine assay on serum at 8 days and 3 months post-TBI

| **Cytokine** | **Treatment Groups** | **p-value** | **Mean difference** | **95% CI of difference** | **DF** | **Sample size** |
| --- | --- | --- | --- | --- | --- | --- |
| **BAFF (8D)** | Sham vs TBI | 0.2590 | 167.1 | -75.47 to 409.6 | 28 | 8, 8 |
|  | TBI vs TBI+NTX | 0.9726 | -38.32 | -280.8 to 204.2 | 28 | 8, 8 |
| **BAFF (3M)** | Sham vs TBI | 0.0340* | 385.9 | 22.88 to 748.9 | 28 | 8, 8 |
|  | TBI vs TBI+NTX | 0.2743 | -245.3 | -608.3 to 117.7 | 28 | 8, 8 |
| **BTC (8D)** | Sham vs TBI | 0.4933 | 1.117 | -1.128 to 3.361 | 14 | 6, 6 |
|  | TBI vs TBI+NTX | >0.9999 | 0.01667 | -2.732 to 2.765 | 14 | 6, 3 |
| **BTC (3M)** | Sham vs TBI | 0.5915 | 1.074 | -1.354 to 3.502 | 15 | 5, 7 |
|  | TBI vs TBI+NTX | 0.5494 | -1.214 | -3.813 to 1.385 | 15 | 7, 4 |
| **Eotaxin (8D)** | Sham vs TBI | 0.0030** | -26.96 | -45.86 to -8.063 | 28 | 8, 8 |
|  | TBI vs TBI+NTX | 0.5363 | 9.388 | -9.512 to 28.29 | 28 | 8, 8 |
| **Eotaxin (3M)** | Sham vs TBI | 0.0033** | -19.03 | -32.52 to -5.531 | 28 | 8, 8 |
|  | TBI vs TBI+NTX | 0.0024** | -19.64 | -33.13 to -6.144 | 28 | 8, 8 |
| **IL-2Ra (8D)** | Sham vs TBI | 0.0231* | -43.95 | -83.03 to -4.870 | 27 | 8, 8 |
|  | TBI vs TBI+NTX | 0.7089 | -15.85 | -56.30 to 24.60 | 27 | 8, 7 |
| **IL-2Ra (3M)** | Sham vs TBI | 0.0024** | -57.44 | -96.75 to -18.13 | 27 | 8, 8 |
|  | TBI vs TBI+NTX | 0.9508 | 7.625 | -31.69 to 46.94 | 27 | 8, 8 |
| **IL-28 (8D)** | Sham vs TBI | 0.1512 | 4.817 | -1.271 to 10.90 | 18 | 8, 6 |
|  | TBI vs TBI+NTX | >0.9999 | -0.01667 | -7.987 to 7.954 | 18 | 6, 3 |
| **IL-28 (3M)** | Sham vs TBI | 0.9853 | -0.5625 | -5.109 to 3.984 | 20 | 8, 8 |
|  | TBI vs TBI+NTX | 0.6272 | -2.125 | -7.036 to 2.786 | 20 | 8, 6 |
| **IL-33 (8D)** | Sham vs TBI | 0.9134 | 4.533 | -14.75 to 23.82 | 22 | 6, 7 |
|  | TBI vs TBI+NTX | 0.9977 | 1.340 | -18.96 to 21.64 | 22 | 7, 5 |
| **IL-33 (3M)** | Sham vs TBI | 0.5378 | -4.452 | -13.65 to 4.748 | 19 | 8, 7 |
|  | TBI vs TBI+NTX | 0.9935 | 1.039 | -10.10 to 12.18 | 19 | 7, 4 |
| **CXCL10 (8D)** | Sham vs TBI | 0.6307 | -1084 | -3809 to 1641 | 10 | 4, 6 |
|  | TBI vs TBI+NTX | 0.6993 | 1236 | -2211 to 4682 | 10 | 6, 2 |
| **CXCL10 (3M)** | Sham vs TBI | 0.6267 | -948.7 | -3258 to 1360 | 12 | 3, 5 |
|  | TBI vs TBI+NTX | 0.9818 | -249.6 | -2249 to 1750 | 12 | 5, 5 |
| **CXCL1 (8D)** | Sham vs TBI | 0.7851 | 1.325 | -2.534 to 5.184 | 28 | 8, 8 |
|  | TBI vs TBI+NTX | 0.8557 | 1.125 | -2.734 to 4.984 | 28 | 8, 8 |
| **CXCL1(3M)** | Sham vs TBI | 0.0005*** | 4.050 | 1.645 to 6.455 | 28 | 8, 8 |
|  | TBI vs TBI+NTX | 0.0977 | -2.125 | -4.530 to 0.2795 | 28 | 8, 8 |
| **Leptin (8D)** | Sham vs TBI | 0.0468* | -203.3 | -404.4 to -2.183 | 27 | 8, 8 |
|  | TBI vs TBI+NTX | 0.7627 | -74.44 | -282.6 to 133.7 | 27 | 8, 7 |
| **Leptin (3M)** | Sham vs TBI | 0.5447 | -102.5 | -310.9 to 106.0 | 28 | 8, 8 |
|  | TBI vs TBI+NTX | 0.0333* | 222.3 | 13.83 to 430.7 | 28 | 8, 8 |
| **CXCL5 (8D)** | Sham vs TBI | >0.9999 | -0.3750 | -60.06 to 59.31 | 28 | 8, 8 |
|  | TBI vs TBI+NTX | 0.5037 | 30.83 | -28.86 to 90.51 | 28 | 8, 8 |
| **CXCL5 (3M)** | Sham vs TBI | >0.9999 | 0.4250 | -48.49 to 49.34 | 28 | 8, 8 |
|  | TBI vs TBI+NTX | 0.0975 | -43.25 | -92.16 to 5.664 | 28 | 8, 8 |
| **CCL7 (8D)** | Sham vs TBI | 0.0882 | 9.425 | -1.021 to 19.87 | 28 | 8, 8 |
|  | TBI vs TBI+NTX | 0.4474 | 5.763 | -4.684 to 16.21 | 28 | 8, 8 |
| **CCL7 (3M)** | Sham vs TBI | <0.0001**** | 51.89 | 30.73 to 73.04 | 28 | 8, 8 |
|  | TBI vs TBI+NTX | 0.5833 | -9.913 | -31.07 to 11.24 | 28 | 8, 8 |
| **RANK L (8D)** | Sham vs TBI | 0.9981 | -5.525 | -92.63 to 81.58 | 26 | 8, 8 |
|  | TBI vs TBI+NTX | 0.7655 | 32.00 | -58.16 to 122.2 | 26 | 8, 7 |
| **RANK L (3M)** | Sham vs TBI | 0.0072** | -64.27 | -113.5 to -15.02 | 25 | 6, 8 |
|  | TBI vs TBI+NTX | 0.0055** | 63.52 | 16.33 to 110.7 | 25 | 8, 7 |
| **GM-CSF (8D)** | Sham vs TBI | 0.9262 | -3.625 | -19.70 to 12.45 | 28 | 8, 8 |
|  | TBI vs TBI+NTX | 0.4711 | -8.625 | -24.70 to 7.447 | 28 | 8, 8 |
| **GM-CSF (3M)** | Sham vs TBI | 0.6175 | -7.193 | -23.35 to 8.962 | 25 | 7, 8 |
|  | TBI vs TBI+NTX | 0.0732 | 15.09 | -1.062 to 31.25 | 25 | 8, 7 |
| **IFN-γ (8D)** | Sham vs TBI | 0.0089** | -36.10 | -64.56 to -7.643 | 28 | 8, 8 |
|  | TBI vs TBI+NTX | 0.9670 | 4.800 | -23.66 to 33.26 | 28 | 8, 8 |
| **IFN-γ (3M)** | Sham vs TBI | 0.1956 | -44.71 | -104.6 to 15.15 | 25 | 8, 7 |
|  | TBI vs TBI+NTX | 0.4671 | 33.14 | -28.68 to 94.96 | 25 | 7, 7 |
| **IL-1β (8D)** | Sham vs TBI | 0.9463 | -9.825 | -59.66 to 40.01 | 22 | 8, 8 |
|  | TBI vs TBI+NTX | 0.4636 | -28.77 | -82.60 to 25.06 | 22 | 8, 6 |
| **IL-1β (3M)** | Sham vs TBI | 0.8511 | 21.00 | -51.40 to 93.14 | 23 | 8, 8 |
|  | TBI vs TBI+NTX | 0.9057 | -18.99 | -96.91 to 58.93 | 23 | 8, 6 |
| **IL-2 (8D)** | Sham vs TBI | 0.8496 | -1.838 | -8.068 to 4.393 | 26 | 8, 8 |
|  | TBI vs TBI+NTX | 0.9964 | -0.5071 | -6.956 to 5.942 | 26 | 8, 7 |
| **IL-2 (3M)** | Sham vs TBI | 0.8777 | 1.735 | -4.686 to 8.156 | 24 | 5, 8 |
|  | TBI vs TBI+NTX | 0.2478 | 3.913 | -1.719 to 9.544 | 24 | 8, 8 |
| **IL-3 (8D)** | Sham vs TBI | 0.9998 | -0.1875 | -6.083 to 5.708 | 28 | 8, 8 |
|  | TBI vs TBI+NTX | >0.9999 | 0.08750 | -5.808 to 5.983 | 28 | 8, 8 |
| **IL-3 (3M)** | Sham vs TBI | 0.3039 | -2.875 | -7.292 to 1.542 | 27 | 8, 8 |
|  | TBI vs TBI+NTX | 0.2463 | 3.196 | -1.375 to 7.768 | 27 | 8, 7 |
| **IL-5 (8D)** | Sham vs TBI | 0.9263 | 6.425 | -22.08 to 34.93 | 28 | 8, 8 |
|  | TBI vs TBI+NTX | 0.9789 | -4.113 | -32.62 to 24.39 | 28 | 8, 8 |
| **IL-5 (3M)** | Sham vs TBI | 0.9404 | -12.89 | -75.02 to 49.25 | 26 | 8, 8 |
|  | TBI vs TBI+NTX | 0.6949 | -25.70 | -90.02 to 38.62 | 26 | 8, 7 |
| **IL-9 (8D)** | Sham vs TBI | 0.9373 | -5.375 | -30.68 to 19.93 | 28 | 8, 8 |
|  | TBI vs TBI+NTX | 0.9830 | 3.388 | -21.92 to 28.70 | 28 | 8, 8 |
| **IL-9 (3M)** | Sham vs TBI | 0.3241 | -18.10 | -46.63 to 10.43 | 26 | 8, 8 |
|  | TBI vs TBI+NTX | 0.9948 | -2.620 | -32.15 to 25.91 | 26 | 8, 7 |
| **IL-10 (8D)** | Sham vs TBI | 0.3441 | 32.95 | -20.14 to 86.04 | 27 | 7, 8 |
|  | TBI vs TBI+NTX | 0.5381 | -25.38 | -76.67 to 25.92 | 27 | 8, 8 |
| **IL-10 (3M)** | Sham vs TBI | 0.6895 | -19.45 | -67.98 to 29.08 | 24 | 8, 8 |
|  | TBI vs TBI+NTX | 0.0162* | -59.45 | -109.7 to -9.220 | 24 | 8, 7 |
| **IL-13 (8D)** | Sham vs TBI | 0.6290 | 11.25 | -16.60 to 39.63 | 12 | 6, 2 |
|  | TBI vs TBI+NTX | 0.9885 | -3.070 | -31.88 to 25.74 | 12 | 2, 5 |
| **IL-13 (3M)** | Sham vs TBI | 0.5779 | -8.025 | -26.51 to 10.46 | 11 | 4, 6 |
|  | TBI vs TBI+NTX | 0.9775 | 2.683 | -17.57 to 22.94 | 11 | 6, 3 |
| **IL-17 (8D)** | Sham vs TBI | 0.3626 | -3.413 | -9.022 to 2.197 | 28 | 8, 8 |
|  | TBI vs TBI+NTX | 0.9448 | 1.138 | -4.472 to 6.747 | 28 | 8, 8 |
| **IL-17 (3M)** | Sham vs TBI | 0.9791 | -1.100 | -8.797 to 6.597 | 26 | 8, 8 |
|  | TBI vs TBI+NTX | 0.5706 | -3.650 | -11.35 to 4.047 | 26 | 8, 8 |
| **CCL2 (8D)** | Sham vs TBI | 0.2548 | 11.39 | -5.056 to 27.83 | 28 | 8, 8 |
|  | TBI vs TBI+NTX | 0.7678 | -5.838 | -22.28 to 10.61 | 28 | 8, 8 |
| **CCL2 (3M)** | Sham vs TBI | 0.9819 | 2.250 | -14.27 to 18.77 | 27 | 8, 8 |
|  | TBI vs TBI+NTX | 0.2753 | -11.52 | -28.62 to 5.573 | 27 | 8, 7 |
| **M-CSF (8D)** | Sham vs TBI | 0.0074** | 3.263 | 0.7451 to 5.780 | 27 | 8, 8 |
|  | TBI vs TBI+NTX | 0.8508 | -0.7679 | -3.374 to 1.838 | 27 | 8, 7 |
| **M-CSF (3M)** | Sham vs TBI | 0.4855 | -1.138 | -3.293 to 1.018 | 28 | 8, 8 |
|  | TBI vs TBI+NTX | 0.7520 | -0.7875 | -2.943 to 1.368 | 28 | 8, 8 |
| **VEGF (8D)** | Sham vs TBI | 0.0397* | 77.90 | 2.847 to 153.0 | 28 | 8, 8 |
|  | TBI vs TBI+NTX | 0.2347 | -53.34 | -128.4 to 21.72 | 28 | 8, 8 |
| **VEGF (3M)** | Sham vs TBI | 0.6103 | 47.38 | -57.33 to 152.1 | 28 | 8, 8 |
|  | TBI vs TBI+NTX | 0.2815 | -70.14 | -174.8 to 34.57 | 28 | 8, 8 |
| **IL-1α (8D)** | Sham vs TBI | 0.0018** | -0.8262 | -1.370 to -0.2825 | 23 | 6, 7 |
|  | TBI vs TBI+NTX | 0.0061** | 0.7262 | 0.1825 to 1.270 | 23 | 7, 6 |
| **IL-1α (3M)** | Sham vs TBI | 0.7785 | 0.1762 | -0.3340 to 0.6864 | 25 | 6, 7 |
|  | TBI vs TBI+NTX | 0.9773 | 0.06964 | -0.4050 to 0.5443 | 25 | 7, 8 |
| **IL-6 (8D)** | Sham vs TBI | 0.0431* | -33.96 | -67.11 to -0.8178 | 28 | 8, 8 |
|  | TBI vs TBI+NTX | 0.0537 | 32.75 | -0.3947 to 65.89 | 28 | 8, 8 |
| **IL-6 (3M)** | Sham vs TBI | <0.0001**** | -66.89 | -100.3 to -33.52 | 27 | 8, 8 |
|  | TBI vs TBI+NTX | 0.3394 | 20.83 | -12.55 to 54.20 | 27 | 8, 8 |
| **IL-12p70 (8D)** | Sham vs TBI | 0.4049 | 34.41 | -24.99 to 93.81 | 28 | 8, 8 |
|  | TBI vs TBI+NTX | 0.2759 | -40.06 | -99.46 to 19.34 | 28 | 8, 8 |
| **IL-12p70 (3M)** | Sham vs TBI | 0.0002*** | -105.8 | -163.8 to -47.81 | 27 | 8, 8 |
|  | TBI vs TBI+NTX | 0.0002*** | 105.0 | 47.02 to 163.0 | 27 | 8, 8 |
| **IL-18 (8D)** | Sham vs TBI | 0.3278 | -3.516 | -9.083 to 2.051 | 26 | 7, 8 |
|  | TBI vs TBI+NTX | 0.0390* | 5.600 | 0.2219 to 10.98 | 26 | 8, 8 |
| **IL-18 (3M)** | Sham vs TBI | >0.9999 | -0.1875 | -8.872 to 8.497 | 28 | 8, 8 |
|  | TBI vs TBI+NTX | 0.2600 | -5.975 | -14.66 to 2.710 | 28 | 8, 8 |
| **IL-23 (8D)** | Sham vs TBI | 0.2940 | -253.9 | -640.4 to 132.7 | 25 | 8, 8 |
|  | TBI vs TBI+NTX | 0.0189* | 510.3 | 69.49 to 951.0 | 25 | 8, 5 |
| **IL-23 (3M)** | Sham vs TBI | 0.7223 | -163.8 | -589.9 to 262.4 | 28 | 8, 8 |
|  | TBI vs TBI+NTX | 0.9341 | 92.16 | -334.0 to 518.3 | 28 | 8, 8 |
| **TNFα (8D)** | Sham vs TBI | 0.0012** | -115.4 | -189.2 to -41.48 | 27 | 7, 8 |
|  | TBI vs TBI+NTX | 0.0104* | 89.00 | 17.64 to 160.4 | 27 | 8, 8 |
| **TNFα (3M)** | Sham vs TBI | 0.8478 | 34.59 | -82.82 to 152.0 | 24 | 6, 8 |
|  | TBI vs TBI+NTX | 0.9750 | 17.01 | -95.51 to 129.5 | 24 | 8, 7 |
| **CCL5 (8D)** | Sham vs TBI | <0.0001**** | -5.425 | -8.258 to -2.592 | 28 | 8, 8 |
|  | TBI vs TBI+NTX | 0.0014** | 4.325 | 1.492 to 7.158 | 28 | 8, 8 |
| **CCL5 (3M)** | Sham vs TBI | 0.7865 | -1.000 | -3.921 to 1.921 | 28 | 8, 8 |
|  | TBI vs TBI+NTX | 0.4876 | -1.538 | -4.458 to 1.383 | 28 | 8, 8 |
| **IL-4 (8D)** | Sham vs TBI | 0.0239* | 2.668 | 0.2864 to 5.049 | 26 | 8, 7 |
|  | TBI vs TBI+NTX | 0.0095** | -3.005 | -5.387 to -0.6239 | 26 | 7, 8 |
| **IL-4 (3M)** | Sham vs TBI | 0.8744 | 0.6036 | -1.614 to 2.821 | 23 | 7, 8 |
|  | TBI vs TBI+NTX | 0.9406 | 0.4750 | -1.839 to 2.789 | 23 | 8, 6 |

Multiplex custom cytokine array on serum samples. A total of 32 analytes, including proinflammatory and anti-inflammatory cytokines and chemokines, and growth factors were investigated during the acute and chronic phases of injury. *p<0.05, **p<0.01, ***p<0.001, ****p<0.0001 with respect to sham vs TBI, and TBI vs TBI/NTX. One-way ANOVA with Tukey’s post-hoc analysis. Abbreviations: TBI, traumatic brain injury; NTX, naltrexone. For methods, see Supporting Information 5.

**Supporting Table 3.** Quantification of NeuN positive cells in the neocortex

|  | **Treatment Groups** | **p-value** | **Mean difference** | **95% CI of difference** | **DF** | **Sample size** |
| --- | --- | --- | --- | --- | --- | --- |
| **8 days** | Sham vs TBI | 0.0150* | 526.9 | 88.73 to 965.0 | 20 | 6, 6 |
|  | TBI vs TBI+NTX | 0.0165* | -520.1 | -958.2 to -81.98 | 20 | 6, 6 |
| **3 months** | Sham vs TBI | 0.9913 | 39.94 | -344.0 to 423.9 | 21 | 6, 6 |
|  | TBI vs TBI+NTX | 0.9772 | 53.55 | -316.4 to 423.5 | 21 | 6, 7 |

Comparison between NeuN (neuronal markers) positive cells between the groups at 8 days and 3 months post-TBI. A significant neuronal loss was observed in the regions with high FJB positive cells in the cortex. NTX prevented the cell loss. *p<0.05 for sham vs TBI, and TBI vs TBI+NTX. One-way ANOVA with Tukey’s post-hoc analysis.

**Supporting Table 4.** Fractional anisotropy analyses from DTI-MRI

| Region | **OB** | **GP** | **HYP** | **CBL** | **FIM** | **LV** | **3V** | **AC** | **MBU** |
| --- | --- | --- | --- | --- | --- | --- | --- | --- | --- |
| Groups | **Pre Post** | **Pre Post** | **Pre Post** | **Pre Post** | **Pre Post** | **Pre Post** | **Pre Post** | **Pre Post** | **Pre Post** |
| TBI (8 d) | 0.259 0.233* | 0.409 0.382* | 0.342 0.322* | 0.291 0.265* | 0.482 0.440* | 0.314 0.263* | 0.234 0.222* | 0.368 0.331* | 0.352 0.342* |
| TBI+NTX (8 d) | 0.268 0.265 | 0.449 0.453 | 0.323 0.331 | 0.302 0.304 | 0.445 0.431 | 0.280 0.290 | 0.228 0.236 | 0.358 0.363 | 0.367 0.354 |
| TBI (3 M) | 0.277 0.245* | 0.365 0.330 | 0.338 0.295 | 0.269 0.242* | 0.481 0.398* | 0.338 0.313* | 0.240 0.237 | 0.362 0.350* | 0.339 0.300* |
| TBI+NTX (3 M) | 0.268 0.243 | 0.449 0.388* | 0.323 0.278* | 0.302 0.253* | 0.445 0.385* | 0.280 0.236 | 0.228 0.213 | 0.358 0.318 | 0.367 0.320 |

**Supporting Table 4 continued.** Fractional anisotropy analyses from DTI-MRI

| Region | **FX** | **4V** | **HBU** | **SPC** | **PG** | **AMY** | **MED** | **PON** |
| --- | --- | --- | --- | --- | --- | --- | --- | --- |
| Groups | **Pre Post** | **Pre Post** | **Pre Post** | **Pre Post** | **Pre Post** | **Pre Post** | **Pre Post** | **Pre Post** |
| TBI (8 d) | 0.342 0.331* | 0.282 0.271* | 0.343 0.365 | 0.139 0.123* | 0.265 0.234 | 0.439 0.439* | 0.333 0.319* | 0.359 0.332* |
| TBI+NTX (8 d) | 0.316 0.316 | 0.276 0.285 | 0.393 0.380 | 0.024 0.025 | 0.266 0.256 | 0.448 0.427 | 0.363 0.360 | 0.408 0.375 |
| TBI (3 M) | 0.347 0.299* | 0.282 0.231* | 0.314 0.259* | 0.254 0.229* | 0.284 0.292 | 0.410 0.392 | 0.334 0.291* | 0.371 0.333* |
| TBI+NTX (3 M) | 0.316 0.276 | 0.276 0.227* | 0.393 0.331* | 0.024 0.029 | 0.266 0.226 | 0.448 0.383* | 0.363 0.316 | 0.408 0.342 |

Mean FA values of different brain regions between TBI and TBI+NTX groups at 8 days and 3 months. All the post-injury values were normalized against the pre-injury group. Therefore, the higher the value, the higher is the FA, whereas lower values represent decreased FA. The decrease in FA was higher in TBI at both 8 days and 3 months, whereas lesser or no changes in FA were observed in the TBI+NTX group, compared to the pre-injury group. An Asterisk represents a decrease in FA after injury. Abbreviations: Olfactory Bulb (OB), Globus Pallidus (GP), Hypothalamus (HYP), Cerebellum (CBL), Fimbria (FIM), Lateral Ventricles (LV), 3rd Ventricle (3V), Anterior Commissure (AC), Midbrain Unsegmented (MBU), Fornix (FX), 4th Ventricle (4V), Hind Brain Unsegmented (HBU), Spinal Cord (SPC), Pineal Gland (PG), Amygdala (AMY), Medulla (MED), Pons (PON)

**Supporting Figure 1.** Standardization of the Experimental Design

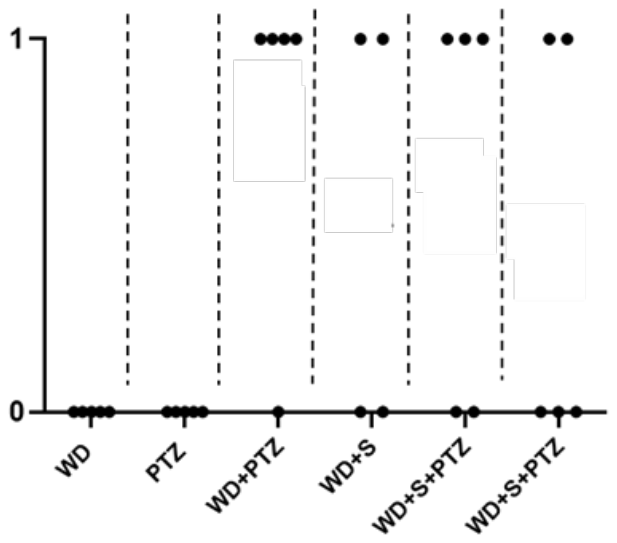

| **Group 🡪** | **WD**  **(Grp I)** | **PTZ**  **(Grp II)** | **WD+PTZ**  **(Grp III)** | **WD+S**  **(Grp IV)** | **WD+S+PTZ**  **Grp (V)** | **WD+S+PTZ**  **(Grp VI)** |
| --- | --- | --- | --- | --- | --- | --- |
| **Number of Animals** | 0 | 0 | 0 | 1 | 1 | 0 |
|  | 0 | 0 | 1 | 0 | 0 | 0 |
|  | 0 | 0 | 1 | 0 | 1 | 1 |
|  | 0 | 0 | 1 | 1 | 0 | 0 |
|  | 0 | 0 | 1 | **-** | 1 | 1 |

Pilot experiments were performed in six different groups (n=4-5) to check mortality, prior to the main experiment. Traumatic brain injury induced by WD (Grp I) or PTZ (Grp II) on its own did not cause any mortality, whereas PTZ, when administered on the same day of WD injury, caused 80% mortality (Grp III). In group IV, about half of the animals died, when surgery was performed after WD on the same day. In group V and VI, TBI was induced on day 1, electrodes were implanted on day 2 and the PTZ test was performed on day 3 and 4. 3 out 5 animals in group V and 2 out 5 animals on day 4 died after the PTZ test. These observations from the pilot studies, and other supporting data, were used to design the main experiment shown in supporting figure 2. Abbreviations: TBI, traumatic brain injury; WD, weight drop; PTZ, pentylenetetrazol.

**Supporting Figure 2.** Neurobehavioral status of the mice after TBI

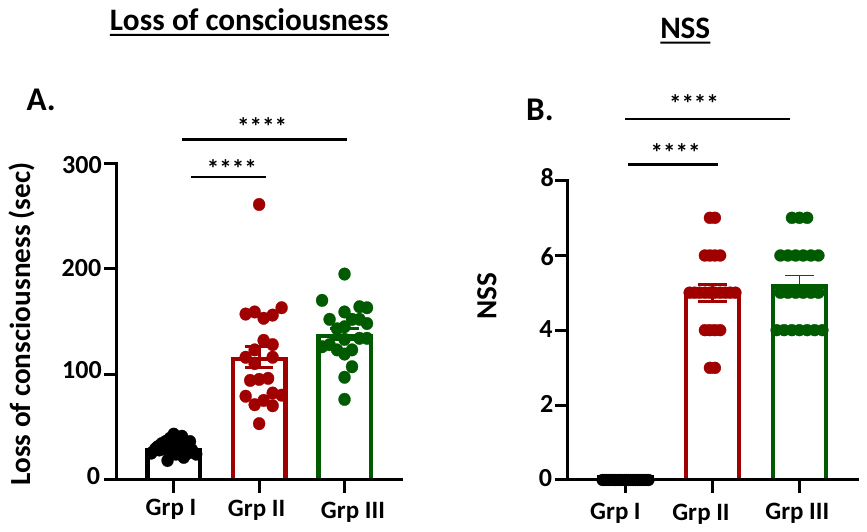

**Functional outcome after TBI.** The severity of TBI was determined by two factors, ie., LOC and NSS. A) LOC is the time time taken by the animal to regain consciousness after TBI. LOC was significantly higher in all the mice that had TBI (Grp II and Grp III), when compared to sham (Grp I). Animals in Grp II and Grp III, after TBI, were randomly selected for the TBI and TBI+NTX groups. The NSS test consists of assessments of various individual parameters which include tests for reflexs (forelimb and hindlimb, pinna, cornea, eye blink and startle reflex tests), inability to walk (hemiplegia, monoparesis, hemiparesis), inability to exit, loss of seeking behavior, prostation, whisking and tremors. If a loss or impairment in any of these behaviors was observed, then a value of 1 was assigned to the animal. Therefore, higher values indicate severe neurological impairment. The experiment was repeated twice with each animal ending up with a total of three injuries (one injury/hour). ****p < 0.0001, One-way ANOVA with Tukey's post-hoc test; *n* = 22. Abbreviations: TBI, traumatic brain injury; LOC, loss of consciousness; NSS, neurological severity score.

**Raw Western Blots**

**Western Blot 1.** S375

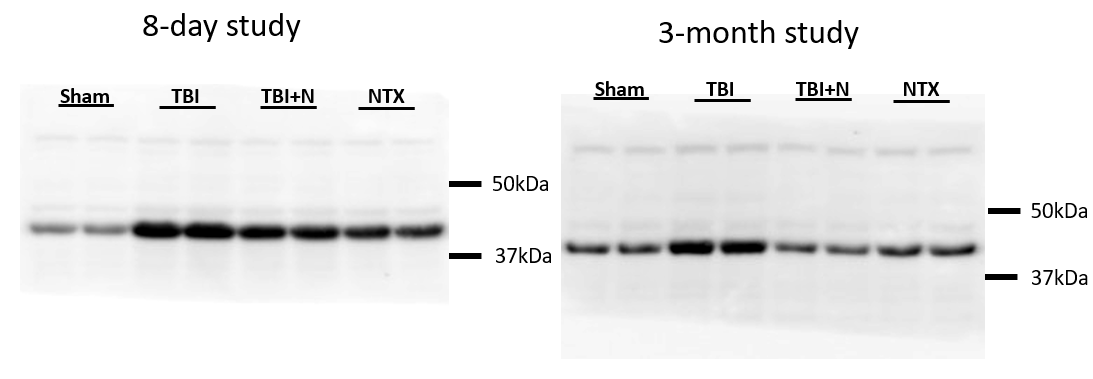

**Western Blot 2.** MOR

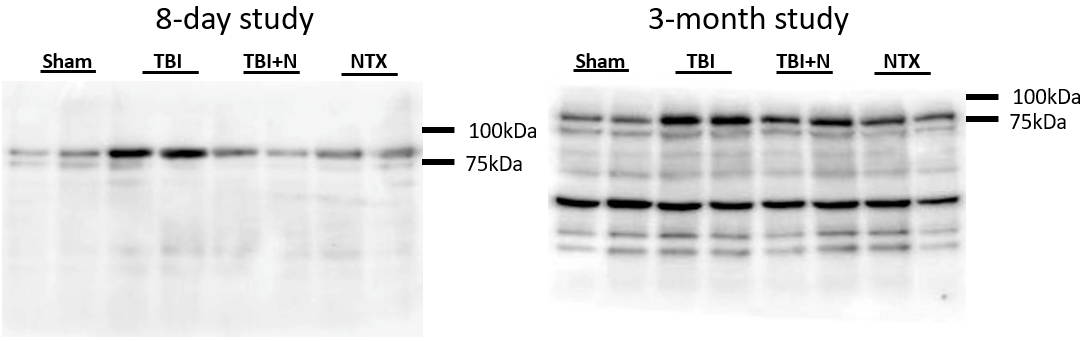

**Western Blot 3.** Phospho-p38

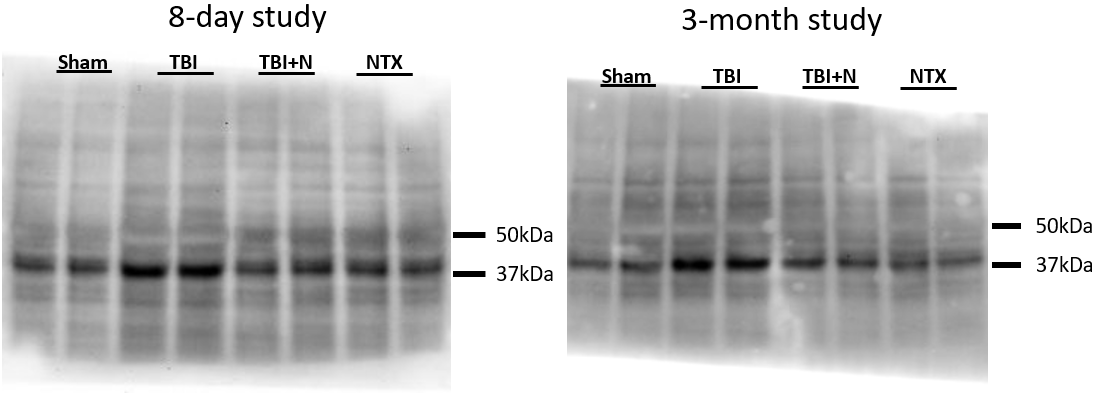

**Western Blot 4.** 3-NT

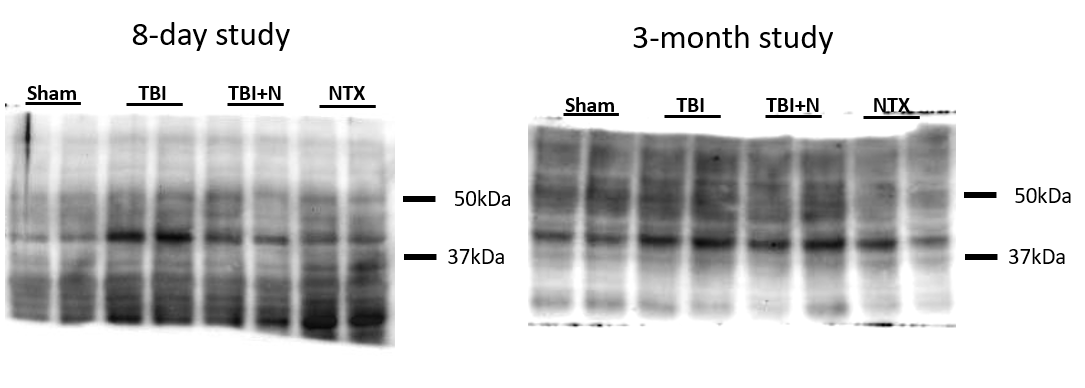

**Western Blot 5.** iNOS

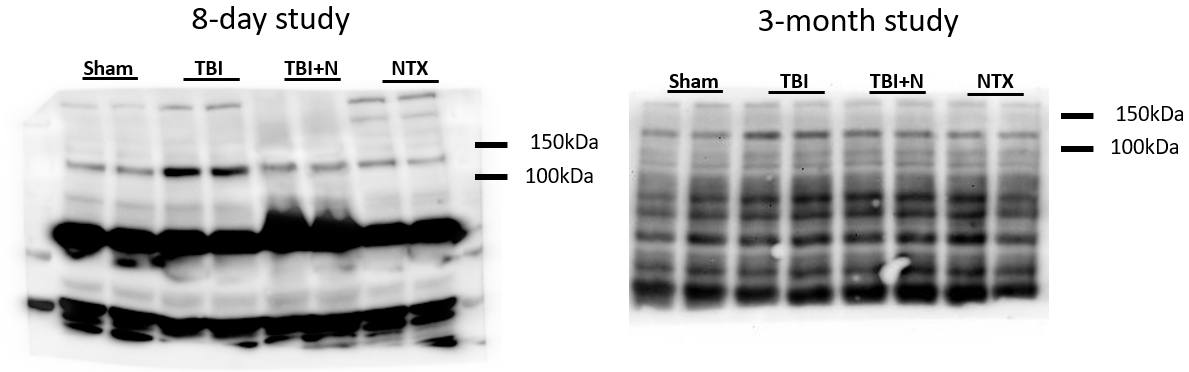
